# Structure and function of the nairovirus cap-snatching endonuclease

**DOI:** 10.1101/2025.10.28.684947

**Authors:** Wenhua Kuang, Zhenhua Tian, Gan Zhang, Fan Wu, Jinyue Li, Jingjing Tang, Huanyu Zhang, Xinyue Zhuo, Zhihong Hu, Manli Wang, Haiyan Zhao, Zengqin Deng

## Abstract

Nairoviruses include several human pathogens such as Crimean-Congo hemorrhagic fever virus (CCHFV) and Kasokero virus (KASV). The cap-snatching endonuclease (EN) domain of the viral polymerase is essential for transcription and represents a promising antiviral target. However, the structural and functional mechanisms of nairovirus ENs remain poorly understood. Here, we describe biochemical and structural studies of the ENs from CCHFV and KASV. Biochemical assays demonstrate that the RNA endonuclease activity of both ENs is activated by manganese ions and exhibits a preference for uridine-rich RNA substrates. This activity is inhibited by three metal-chelating inhibitors (DPBA, L-742,001, and BXA), with BXA displaying the highest binding affinity and inhibitory potency. We further determine nine crystal structures of CCHFV and KASV ENs in apo, metal ion-bound, and inhibitor-bound states. Comparative structural analysis uncovers a two-metal-ion binding mode unique to nairovirus ENs, in which conserved residues coordinate two manganese ions via bridging water molecules. In the inhibitor-bound structures of KASV EN, BXA forms additional stabilizing interactions with the enzyme, explaining its superior inhibitory effect. Functional assays further confirm that the two-metal-ion mechanism is critical for viral transcription. These findings provide a structural foundation for the rational design of antivirals against CCHFV and related pathogens.

## Introduction

Nairoviruses are a large group of tick-borne bunyaviruses, including several highly pathogenic agents infecting humans and animals. Among them, the Crimean-Congo hemorrhagic fever virus (CCHFV) is the most important human pathogen. CCHFV is endemic in over 30 countries across Asia, Europe, and Africa. Infection with CCHFV can cause serious diseases such as acute fever, hemorrhagic symptoms, and multi-organ failure, with a case fatality rate of up to 40% (1,2). Due to its high infectivity, pathogenicity, and the lack of approved vaccines and antiviral drugs, CCHFV is classified as a Biosafety Level 4 (BSL-4) pathogen and has been listed by the World Health Organization (WHO) as a Priority Pathogen requiring urgent research to develop countermeasures (3,4). Infections with other members of nairoviruses, Kasokero virus (KASV), Dugbe virus (DUGV), and Erve virus (EREV), can also cause symptoms in humans, including headache, rash, diarrhea, thrombocytopenia, and neurological disorders (5–7). In livestock, infection of Nairobi sheep disease virus (NSDV) causes hemorrhagic gastroenteritis. In recent years, an increasing number of emerging nairoviruses have been identified (8–10), raising concerns about their potential biosecurity risks to public health.

Segmented negative-strand RNA viruses (sNSVs), including bunyaviruses and orthomyxoviruses, employ a cap-snatching mechanism to initiate transcription of their viral genomes. This process is mediated by the N-terminal endonuclease (EN) domain of RNA-dependent RNA polymerase (RdRP, also known as L protein in bunyaviruses), which cleaves host-derived capped RNA fragments to serve as primers for viral mRNA synthesis (11,12). The cap-snatching EN domain is a metal ion-dependent nuclease characterized by a conserved PD… D/ExK catalytical motif. Structural and biochemical studies have classified viral ENs into two groups, His+ ENs and His− ENs, based on their active site architecture and enzymatic activity (13) (**Figure 1A**). His+ ENs, found in the families of *Peribunyaviridae*, *Phenuiviridae*, and *Hantaviridae*, and influenza viruses, contain a conserved histidine that coordinates the first metal ion, along with a flexible active-site loop harboring an acidic residue that binds the second metal ion (14–19). These enzymes exhibit robust *in vitro* endonuclease activity. By contrast, His− ENs, exclusive to *Arenaviridae*, substitute the histidine with glutamate or aspartate for metal ion coordination and possess a distal flexible loop that does not participate in metal ion coordination (18,20,21). Strikingly, His− ENs demonstrate minimal enzymatic activity in *in vitro* assays.

**Figure 1.**
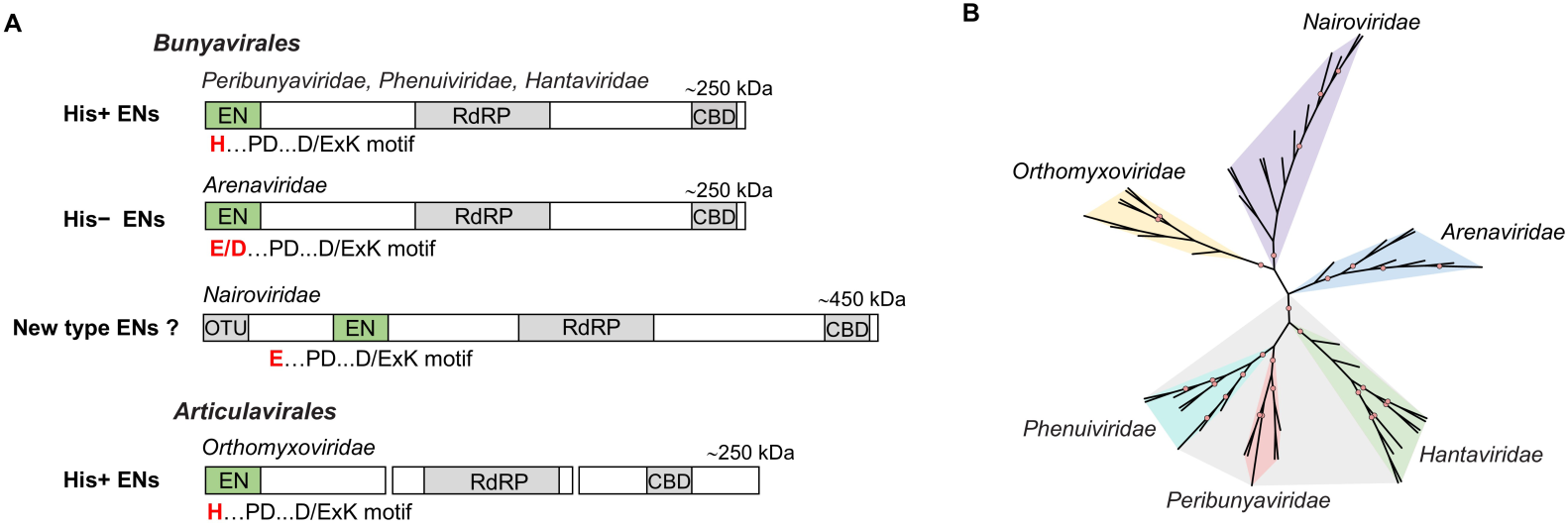
Nairovirus ENs differ from the other two categories of ENs. (A) Schematic representation of the cap-snatching endonuclease domains in the RNA polymerase of sNSVs. Nairovirus ENs show an apparent discrepancy in domain location and active site motifs compared to the known ENs (His+ and His− ENs). Identified functional domains in the RNA polymerase, including endonuclease (EN), RNA-dependent RNA polymerase (RdRP), ovarian tumor domain (OTU), and cap-binding domain (CBD), are indicated. (B) An unrooted phylogenetic tree of representative ENs from five human-infecting bunyavirus families. The sequence boundaries of 51 EN domains from five *Bunyaviridaes* were determined by aligning the L proteins, using the characterized EN domain as the query sequence. Subsequently, the amino acid sequences of EN homologs were aligned using MAFFT (51). Phylogenetic analysis was performed using IQ-TREE (52), with branch support evaluated using SH-like approximate likelihood ratio test (SH-aLRT) and ultrafast bootstrap (UFBoot) with 1000 replicates. Nodes with strong support (SH-aLRT ≥ 80% and UFBoot ≥ 95%) are marked by pink circles in the tree. The UniProt accession numbers and sequence boundaries for the 51 EN domains are listed in Table S2.

Compared to His+ ENs and His− ENs, the nairovirus ENs exhibit a hybrid active-site motif, integrating key features from both classes. Specifically, they retain a conserved glutamate residue, a hallmark of His− ENs, which is responsible for the first metal ion coordination, while also possessing a flexible loop region characteristic of His+ ENs, which is likely involved in the second metal ion binding (22). In addition to these hybrid features, nairovirus ENs differ from other bunyavirus ENs in both domain location and overall composition (**Figure 1A**). While most bunyavirus ENs are located at the N-terminus of the polymerase and comprise about 200 residues, the nairovirus ENs are positioned at nearly 500 residues downstream from the N-terminal and span approximately 350 residues. Moreover, nairovirus ENs contain one or two unique insertions (30-40 residues, insertion 1 and 2), predicted to form highly flexible loops that are absent in other viral ENs. Phylogenetic analysis of bunyaviruses further highlights the distinctiveness of nairovirus ENs. Representative ENs from five human-infecting bunyavirus families cluster into three major clades (Table S2), two of which align with the known His+ ENs and His− ENs (**Figure 1B**). Notably, nairovirus ENs form a separate clade, suggesting evolutionary divergence from these canonical groups. Collectively, these striking distinctions imply that nairovirus EN likely represents a class of viral endonucleases with different enzymatic and structural characteristics. However, the metal ion-dependent catalytic mechanism of nairovirus ENs remains unresolved, primarily due to the lack of high-resolution structural information and limited biochemical characterization of these ENs.

The cap-snatching endonuclease, a universally conserved feature among sNSVs, plays a pivotal role in viral transcription, making it an attractive target for antiviral development. Given the metal ion-dependent activity of ENs, a wealth of metal-chelating inhibitors has been explored to target ENs. Diketo acid compounds represent the first class of inhibitors developed to target the influenza virus EN. Among them, 2,4-dioxo-4-phenylbutanoic acid (DPBA), the prototype of diketo acid compounds, inhibits the EN activity of influenza virus with a half-maximal inhibitory concentration (IC_50_) at tens of micromolar levels (23). Subsequently, a derivative of DPBA, L-742,001, emerged as a more potent compound, exhibiting strong inhibition of both endonuclease activity and cell-based virus replication at submicromolar concentrations (24,25). More recently, baloxavir acid (BXA), has emerged as a highly potent inhibitor, demonstrating nanomolar antiviral activity *in vitro* and therapeutic efficacy in mouse models against influenza virus (26–28). Its prodrug, baloxavir marboxil, has been approved by the FDA as the first clinically available EN-targeting drug for influenza treatment (29). Notably, DPBA, L-742,001, and BXA have also shown varying degrees of inhibitory effects against multiple bunyaviruses in enzymatic and cell-based assays, underscoring their potential as broad-spectrum antivirals due to their conserved metal-chelating pharmacophore. Compared to their potency against influenza virus, these inhibitors demonstrate suboptimal efficacy against bunyaviruses, with IC₅₀ values ranging from several micromolar to several hundred micromolar (30–33), highlighting the need for further optimization. However, the detailed molecular mechanisms underlying the action of these inhibitors remain poorly understood, posing a challenge to rational drug optimization and further antiviral development against bunyaviruses.

Currently, there is a gap in our understanding of the structure and function of nairovirus ENs, which is critical for comprehensively elucidating the catalytical mechanisms of viral ENs and developing effective antiviral strategies against CCHFV and related nairoviruses. In this study, we present the crystal structure of CCHFV EN in apo form and a series of high-resolution crystal structures of KASV EN in apo form, Mn^2+^-bound form, and in complex with three representative EN inhibitors. We biochemically characterized nairovirus ENs’ metal ion-dependent endonuclease activity, investigated the functional roles of conserved active site residues in EN catalysis, and evaluated the inhibitors’ binding affinity and inhibitory activity. Our findings reveal a two-metal-ion binding mode specific to the *Nairoviridae* family, which is essential for achieving their full catalytic activity. Furthermore, we found that the FDA-approved inhibitor BXA exhibits potent inhibitory activity against nairovirus ENs and engages in more extensive interactions with the active site pocket of KASV EN than the other two inhibitors (DPBA and L-742,001). Our combined biochemical, structural, and functional analyses provide important insights into the catalytic and inhibitory mechanisms of nairovirus endonucleases, offering potential opportunities for structure-based drug design and optimization.

## Materials and Methods

### Protein expression and purification

The fragments containing putative endonuclease domain were designed based on bioinformatics analysis. DNA fragments encoding residues 585-898 of CCHFV L protein and 610-904 for KASV L protein were amplified by PCR using synthesized gene sequences and subsequently cloned into the pGEX-6P-1 expression vector with an N-terminal GST tag. Single-point mutations were introduced by site-directed mutagenesis. Proteins were expressed in *E. coli* strain BL21 (DE3) cultured in LB medium supplemented with 100 μg/ml ampicillin at 16°C overnight after induction with 0.2 mM of IPTG. Cells were harvested, resuspended in lysis buffer (20 mM Tris-HCl pH 8.0, 300 mM NaCl), and disrupted using a high-pressure cell crusher (Union-Biotech). The clarified lysate was loaded onto a column packed with Glutathione Resin, washed with 20 column volumes of lysis buffer, and subjected to on-column cleavage with PreScission protease at 4°C overnight to remove the GST tag. The flow-through fraction containing target proteins was concentrated and further purified by size-exclusion chromatography using a Superdex 200 Increase 10/300 GL column (Cytiva) equilibrated with GF buffer (20 mM Tris-HCl pH 8.0, 150 mM NaCl). Purified proteins were concentrated to 10-25 mg/ml, flash-frozen in liquid nitrogen and stored as aliquots at −80°C for further use. Mutant proteins were prepared using the same purification strategy as the wild-type (WT) protein.

### Differential scanning fluorimetry (DSF)

DSF thermal denaturation curves of purified KASV EN and mutants were measured using a PSA-16 instrument (BEST Science & Technology, Beijing, China). Protein samples were diluted to 0.3 mg/mL in assay buffer (20 mM Tris-HCl pH 7.5, 150 mM NaCl, 5 % vol/vol glycerol) supplemented with 2 mM of various divalent metal ions, or with 0.2 mM of inhibitor (DPBA, L-742,001 and BXA) and 2 mM MnCl_2_. Intrinsic protein fluorescence intensity at 330 nm and 350 nm was monitored using a linear temperature ramp from 30°C to 95°C at a heating rate of 1°C per minute. The melting temperature (*T*_m_) was calculated from the first derivative of the F350/F330 fluorescence ratio curve.

### *In vitro* EN activity assay

The 5’-FAM-labeled ssRNA substrates including 19-mer U-rich ssRNA (5’- AUUUUGUUUUUAAUAUUUC-3’), A-rich ssRNA (5’-UAAAAGAAAAAUUAUAAAC-3’), C-rich ssRNA (5’- ACCCCGCCCCCAACACCCA-3’), G-rich ssRNA (5’-AGGGGCGGGGGAAGAGGGA-3’), and 27-mer structured ssRNA (5’- GAUGAUGCUAUCACCGCGCUCGUCGUC-3’) were chemically synthesized (Shanghai Sangon Biotech Co., Ltd; Integrated DNA Technologies). For nuclease activity experiments, 1-4 μM of the EN was incubated with 0.2 µM 5’-FAM-labeled ssRNA substrates at 37°C in assay buffer (5 mM Tris-HCl pH 7.5, 150 mM NaCl, 1 U*/*μL RNasin [Promega]), in the presence or absence of 2 mM divalent metal ions, 20 mM EDTA, or inhibitor at the indicated concentration. Reactions were terminated by adding 2 × stop loading buffer (95% formamide, 20 mM EDTA, 0.02% wt/vol bromophenol blue). Samples were heated at 100°C for 1 min, resolved on 7 M urea, 20% polyacrylamide, Tris-borate-EDTA gel electrophoresis, and then visualized with Typhoon imager (GE Healthcare). The intensity of the ssRNA substrate in the images was quantified using ImageJ, (https://imagej.nih.gov/ij), and the enzymatic activity is determined by calculating the percentage of substrate ssRNA degraded. The activity for the mutants is expressed relative to the WT EN (set at 100%). For activity inhibition assays, inhibition percentages were plotted against compound concentrations, and the dose-response curves were generated by nonlinear regression analysis using GraphPad Prism 9 to determine half-maximal inhibitory concentrations (IC_50_).

### mAb generation

BALB/c mice were immunized three times at three-week intervals with CCHFV or KASV EN proteins. Following the final immunization, mice were euthanized and splenocytes were harvested for antigen-specific B cell sorting and antibody sequence identification. Briefly, isolated cells were stained with 0.5 μg/mL biotinylated EN proteins for 30 minutes at 4°C. After washing with PBS, color conjugated secondary antibodies were added for 30 minutes at 4°C: Dead 780-APC-Cy7, CD3/CD4/CD8-AmCyan, CD19-PE-Cy7, IgD-PerCP-Cy5.5, CD138-FITC, CD95-PE, and Streptavidin-APC (BD Pharmingen). After two additional washes, cells were sorted on a FACS Aria II (BD Biosciences) using the following gating strategy: Dead-, CD3/CD4/CD8-, CD19+, IgD-, CD95+, CD138+ and Streptavidin-APC+ (EN-specific binding). Target cells were sorted into 96-well PCR plates (Bio-Rad), and the variable regions of antibody genes were amplified using established protocols (34,35). The resulting sequences were cloned into mammalian expression vectors containing the human CH1 domain of the heavy chain and the constant region of the light chain for Fab expression, respectively.

For Fab production, paired heavy- and light-chain plasmids were co-transfected into Expi293 cells at a 1:1 molar ratio. Cell supernatants containing secreted Fabs were harvested 5-6 days post-transfection, filtered through 0.45 μm filters, and purified sequentially using Ni-NTA resin (GenScript, Cat# L00666) followed by size-exclusion chromatography on a Superdex 200 Increase 10/300 GL column equilibrated with GF buffer (20 mM Tris-HCl pH 8.0, 150 mM NaCl).

### Crystallization and structure determination

The purified EN protein was mixed with Fab in a molar ratio of 1:1.2 and further purified to homogeneity via size-exclusion chromatography using Superdex 200 Increase 10/300 GL column. Crystallization trials of EN-Fab complex were conducted using a mosquito robot (TTP Labtech) with the sitting-drop vapor diffusion method at 16°C. Typically, protein samples at concentrations of 8 or 10 mg/mL were mixed with the reservoir solution at a 1:1 volume ratio in 0.6-μl droplets. CCHFV EN-G5 Fab complex crystallized in a reservoir solution containing 2% vol/vol Tacsimate^TM^ pH 7.0, 0.1 M HEPES pH 7.5, and 20% wt/vol Polyethylene glycol 3,350. Crystals of KASV EN-2E9 Fab complex were obtained in the condition of 25% vol/vol Jeffamine ED2003, 0.2 M NaCl, and 0.1 M MES-NaOH pH 6.0. The KASV EN-Mn^2+^ bound crystals were obtained by soaking apo crystals in reservoir solution supplemented with 2.5 mM MnCl_2_ for 6 hours. KASV EN mutant crystals were grown under the same conditions as the WT protein, with the addition of 2.5 mM MnCl_2_. KASV EN-inhibitor complex crystals were obtained by co-crystallization using protein solutions supplemented with 2.5 mM MnCl_2_ and 0.5 mM of individual inhibitor. Apo and mutant crystals were flash-cooled in liquid nitrogen using cryoprotectant solutions consisting of reservoir supplemented with 15% vol/vol glycerol, with or without 2.5 mM MnCl₂. For EN-inhibitor co-crystals, an additional 0.5 mM inhibitor was included in the cryoprotectant.

The X-ray diffraction data were collected at BL02U1 and BL10U2 beam lines of the Shanghai Synchrotron Radiation Facility (SSRF) with a wavelength of 0.979 Å and a temperature of 100 K. A full 360 degrees of data was collected in 0.4 °oscillation steps. The diffraction data were automatically processed at the beamlines using the XDS and Xia2 pipelines (36,37) and subsequently scaled with Aimless from the CCP4 suite (38). The initial model was obtained by the molecular replacement (MR) program PHASER (39) using the predicted EN and Fab structures from AlphaFold2 (40) as search models. Structure refinement was carried out iteratively through manual model building in Coot (41) and automated refinement with PHENIX (42). Data collection and refinement statistics for the final models are summarized in Table S1.

### Microscale thermophoresis (MST) assay

Binding affinities between ENs and inhibitors were measured using the Monolith NT.115 (NanoTemper Technologies). N-terminally GFP-tagged KASV EN and CCHFV EN were prepared and used in the experiments for fluorescence-based detection. In each assay, 10 μl of 200 nM GFP-EN in buffer containing 20 mM Tris-HCl pH 7.5, 150 mM NaCl, 2 mM MnCl_2_, 5% vol/vol DMSO, 0.05% vol/vol Tween-20, 1 mg/mL bovine serum albumin (BSA) was mixed with 10 μl of inhibitor solution (DPBA, L-742,001, or BXA) at various concentrations. After a 15-minute incubation at room temperature, samples were loaded into capillaries (NanoTemper Technologies), and temperature-induced fluorescence changes were recorded at 25°C using 20% LED power and medium MST power. Dissociation constants (Kd) were determined by nonlinear regression analysis of dose-response curves using MO.Affinity Analysis software.

### CCHFV mini-replicon assay

The transcription activities of L protein mutants were assessed using the T7 RNA polymerase-based CCHFV mini-replicon system, as described previously(43). Similarly, 0.5 μg each of the three plasmids pCAGGS-CCHFV-L (WT or mutants), pCAGGS-CCHFV-NP, and pT7-CCHFV-Lutr-eGFP were transfected into 1×10^5^ BSR-T7/5 cells using Lipofectamine 3000 (Invitrogen), following the manufacturer’s instructions. After transfection, cells were incubated at 37°C with 5% CO₂ for 24 hours, then fixed with 4% polyformaldehyde for 15 minutes. Subsequently, the cells were washed three times with PBS and stained with Hoechst 33258 (Beyotime) for 10 minutes to visualize nuclei. The total number of cells and the number of fluorescent cells were counted using a high-content imaging system (HCS; PerkinElmer Operetta CLS, Tokyo, Japan), and the percentage of GFP-expressing cells relative to the total cells was calculated. The ratio of the mutant group was normalized to that of the WT group, which was set as 100%. Statistical analysis was performed using two-tailed Student’s *t* tests.

To confirm the expression of all L protein mutants in BSR-T7/5 cells, transfections were performed using 1 μg of pCAGGS-CCHFV-L plasmid, which expresses C-terminally 3xFLAG-tagged L protein, in a 24-well plate (1×10^5^ cells/well) with Lipofectamine 3000. At 24 hours post transfection, cells were lysed, separated by 8% SDS-PAGE, and transferred to PVDF membrane. L protein was detected using a Mouse anti DDDDK-Tag mAb with a dilution of 1:3,000 (Abclonal, Cat#AE005) and an HRP-conjugated Goat anti-Mouse IgG (H+L) with a dilution of 1:5,000 (Abclonal, Cat#AS003). Protein bands were visualized by ChemiScope 6100 (Clinx Science Instruments Co., Ltd; Shanghai, China).

## Results

### Biochemical and enzymatic characterization of KASV EN

Both characterized His+ and His− cap-snatching endonucleases require divalent metal ions for catalytic activity. To identify the metal ion binding specificity of nairovirus EN, Differential scanning fluorimetry (DSF) was performed to monitor protein stability under varying conditions. KASV EN displays a typical thermal denaturation curve and has a melting temperature (*T*_m_) of 42.3°C in the absence of metal ions. The addition of MnCl₂ or MgCl₂ increased the *T*_m_ by 4.7°C and 2.9°C, respectively (**Figure 2A**), probably due to the metal ions binding in the active site. Abnormal thermal denaturation curves were observed in the presence of Co²⁺, Zn²⁺, or Ni²⁺ (**Figure S5A)**, presumably because these ions lead to protein aggregation or degradation. Further DSF assays were conducted in the presence of MnCl₂ and three known EN inhibitors. Both DPBA and L-742,001 induced an approximately 10°C increase in *T*_m_, surpassing the stabilization effect observed with MnCl₂ alone (Δ*T*_m_ =4.7 °C). Notably, BXA conferred the greatest stabilization effect on KASV EN, increasing the *T*_m_ by 16.8°C (**Figure 2A**). These results indicate that KASV EN can bind both Mn²⁺ and Mg²⁺ and might interact most strongly with BXA among the tested inhibitors.

**Figure 2.**
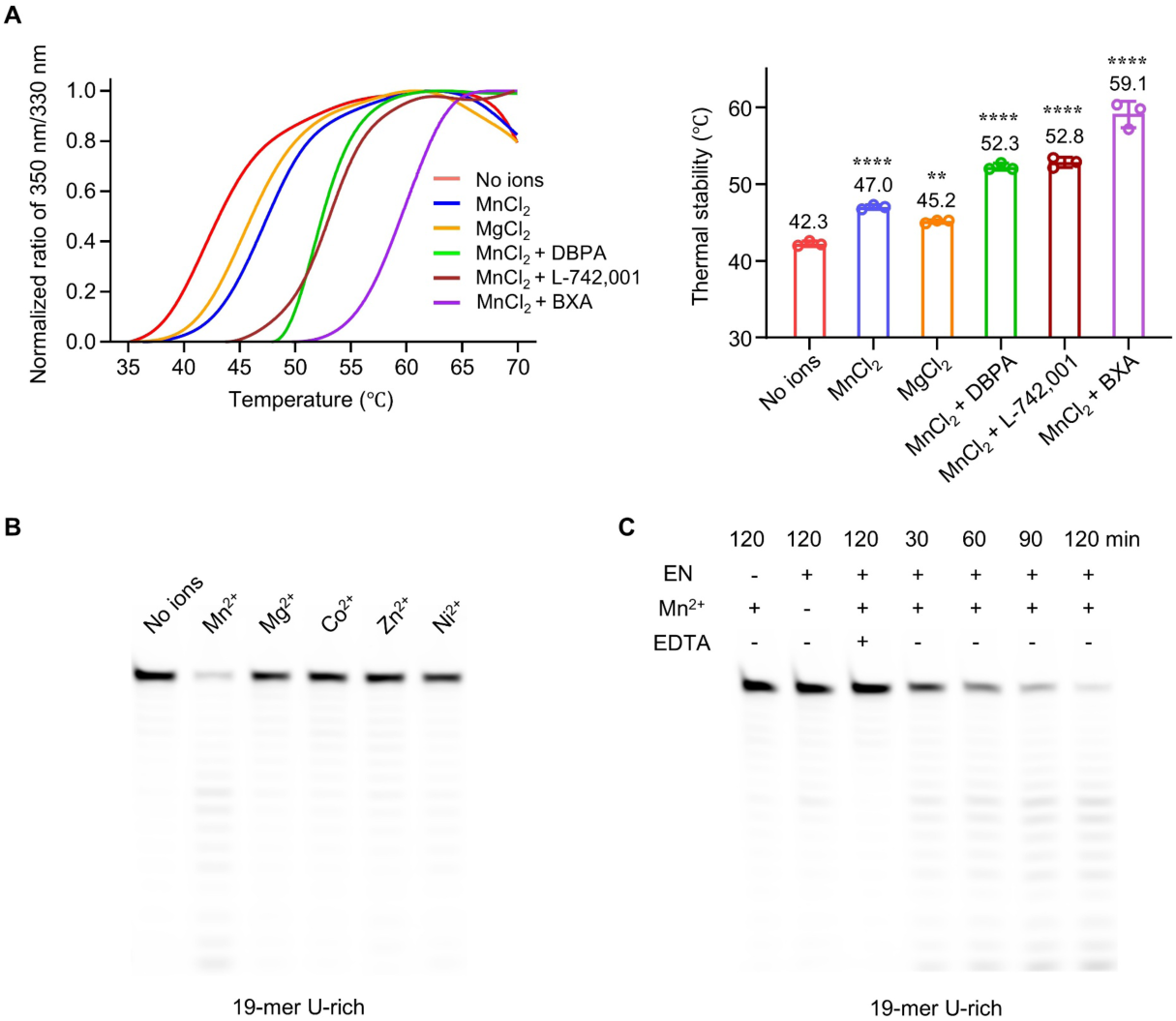
Thermal stability and divalent metal ion-dependent activity of KASV EN. (A) Thermal denaturation profiles of KASV EN in the absence and presence of indicated divalent metal ions and inhibitors. The column chart shows the mean values ± SD of *T*_m_ obtained from three independent experiments. Statistical significance of differences was evaluated via one-way ANOVA analysis. The MnCl₂ and MgCl₂ groups were respectively compared to the no-ion control group, while the inhibitor-treated groups were compared to the MnCl₂ group. \*\**P* < 0.01, \*\*\*\**P* < 0.0001. (B) Divalent metal ion-dependent endonuclease activity of KASV EN. 1 μM of enzyme was incubated with 19-mer U-rich ssRNA for 2 hours at 37℃ in the presence of 2 mM of the indicated metal ions. (C) Time course of endonuclease activity of KASV EN with 2 mM MnCl_2_ and 19-mer U-rich ssRNA as the substrate.

To characterize the intrinsic nuclease activity of KASV EN, we conducted nucleic acid degradation assays with various metal ions and five distinct 5’-FAM-labeled single-stranded RNA (ssRNA) substrates. KASV EN showed apparent nuclease activity only with Mn^2+^ when using the 19-mer U-rich ssRNA, whereas no detectable activity was observed with the other four tested metal ions, confirming a strong specificity for Mn^2+^ (**Figure 2B**). Furthermore, the RNA substrate preference of KASV EN was characterized by comparing the EN activity on 19-mer U-rich, A-rich, C-rich, and G-rich ssRNAs, as well as a 27-mer heterogeneously structured ssRNA. For the nucleotide-rich constructs, identical nucleotide positions were designed to ensure sequence uniformity across substrates. The results showed that the majority of U-rich ssRNA was digested into small fragments by KASV EN within 120 min (**Figure 2C**), whereas the other four ssRNA substrates remained predominantly intact throughout the incubation period (**Figure S1A**). Complete and pronounced degradation was observed for 19-mer U-rich and the other three RNA constructs except for the 19-mer G-rich ssRNA at higher protein concentrations, indicative of KASV EN preference for uridine ssRNAs (**Figure S1B**). We also examined the activity of CCHFV EN under the same experimental conditions, which exhibited analogous RNA digestion profiles as those of KASV EN (**Figure S1B**), suggesting a common substrate preference across nairovirus ENs.

### Antibody-assisted crystallization of CCHFV and KASV ENs

Initial attempts to crystallize the CCHFV and KASV ENs in apo form or in complex with metal ion and inhibitors were unsuccessful, likely due to the inherent conformational flexibility derived from the two insert regions. To address this challenge, we developed an innovative antibody-assisted crystallization strategy. Mice were immunized with EN proteins, and monoclonal antibodies (mAbs) specific to CCHFV EN (G5) and KASV EN (2E9) were successfully identified (**Figure S2A**). Enzymatic assays showed that both mAbs had no effects on the *in vitro* endonuclease activity of CCHFV or KASV ENs (**Figure S2B**). The complex formed between EN and the corresponding antigen-binding fragment (Fab) yielded large rod-like crystals for the CCHFV EN-G5 Fab complex and thin plate-like crystals for the KASV EN-2E9 Fab complex. The CCHFV EN-G5 Fab complex is crystalized with a unit cell containing two complex molecules arranged in a staggered manner with neighboring molecules, which adopt a herringbone packing motif within the crystal lattice. This ordered spatial organization is primarily driven by compact intermolecular interactions between the EN and Fab. The KASV EN-2E9 Fab complex crystal possesses a small-sized unit cell containing a single complex molecule. Within the crystal lattice, two neighboring complexes orient in opposite directions and assemble into extended two-dimensional parallel layers via Fab-Fab interactions (**Figure S2C**). Notably, in both complex structures the Fab binds specifically to flexible loop regions of EN and stabilizes the conformation of individual EN molecule, indicating that Fab-mediated interactions play an indispensable role in crystal packing.

### Overall structures of KASV and CCHFV ENs

The apo and Mn^2+^-bound structures of KASV EN were solved at 1.90 Å and 1.98 Å resolution in space group *P*2_1_, respectively. Clear electron density was visible for the entire protein except for residues 726-779, which coincided with the insertion 1, predicted to be a highly flexible loop. The apo structure of CCHFV EN was solved at 3.05 Å in space group *P*2_1_2_1_2_1_, with two disorder regions corresponding to the insertion 1 (residues 703-729) and insertion 2 (residues 764-795), respectively. The Mn^2+^-bound structure of CCHFV EN could not be obtained due to poor diffraction quality of the complex crystals. Therefore, we focus our detailed analysis on the high-quality KASV EN structures.

Similar to previously reported EN structures, the KASV EN adopts a two-lobe architecture. One lobe includes the N-terminal four helix bundle (α1-3 and α9), and the other is composed of a central five-stranded β-sheet sandwiched by five helices, with the helices α4-6 and α8 partially covering the β-sheet, while helix α7 places at the bottom of the β-sheet (**Figure 3A**). The active site lies between the two lobes and harbors two manganese ions coordinated by highly conserved acidic residues (**Figure S3C)**. The flexible loop linking α3 and α4 (highlighted in green in Figure 3A, denoted as LoopE), bears a conserved acidic residue that is involved in manganese binding, and plays a similar function to that observed in the His+ ENs. The CCHFV EN displays highly consistent global conformations as that of KASV EN, with a root-mean-square deviation (RMSD) value of 1.2 Å for all superimposable *a*-carbon atoms, except for two small helices situated at the two ends of the strand β3 (**Figure 3 and Figure S3A**).

**Figure 3.**
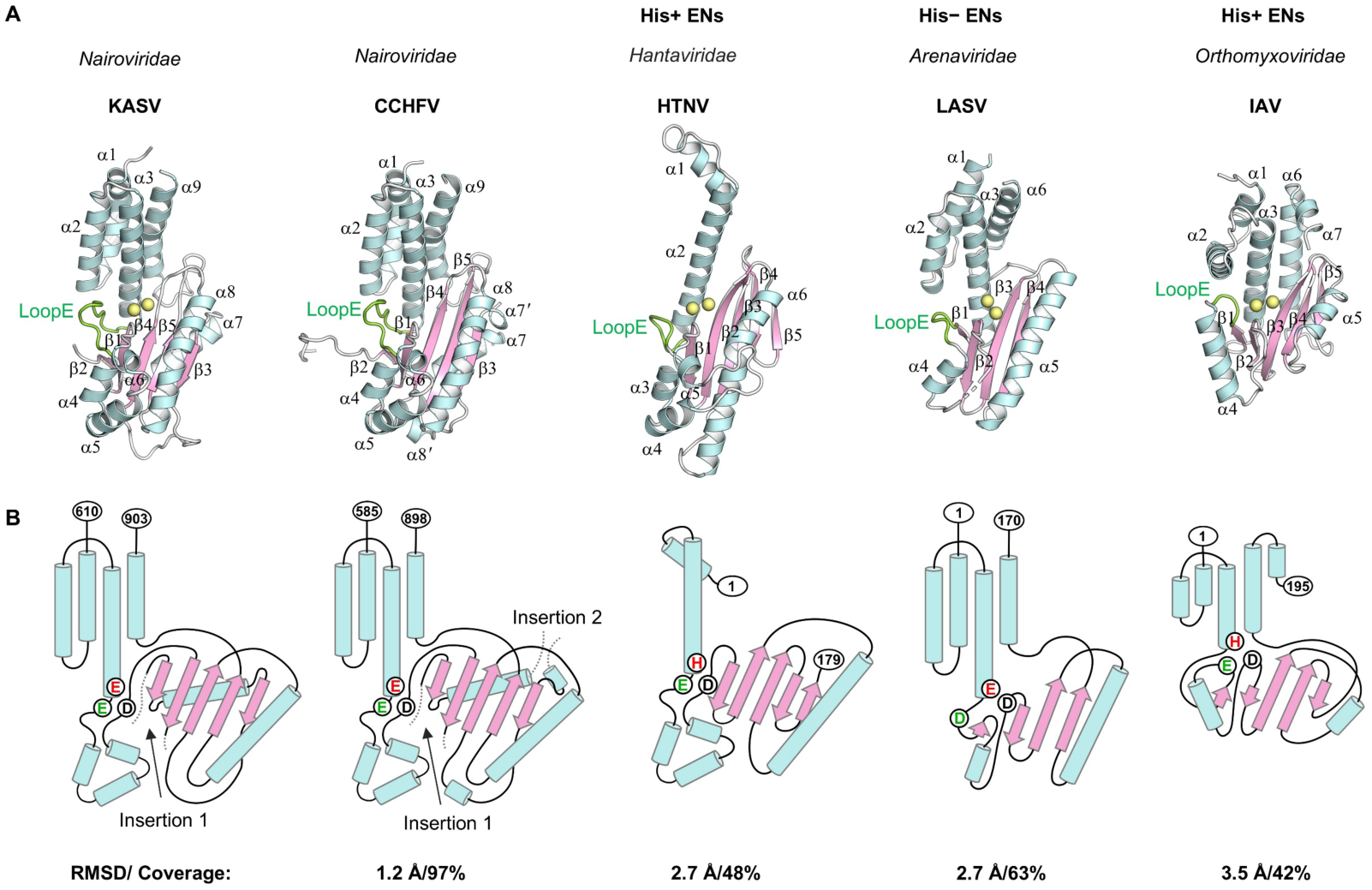
Crystal structures of KASV and CCHFV ENs compared with other sNSV cap-snatching endonucleases. (A) Structural comparison of KASV and CCHFV ENs with His+ and His− endonucleases. The overall structures of ENs are shown in cartoon representation, with α-helices colored in cyan and β-strands in pink. Manganese ions bound in the active site are shown as yellow spheres. The LoopE involved in metal ion coordination is colored green. PDB entries: 5IZE (Hantaan virus, HTNV), 5J1P (Lassa virus, LASV), 7KAF (Influenza A virus, IAV). (B) Topological diagrams of the cap-snatching ENs. The key metal ion-coordination residues are highlighted in red circles. Narovirus ENs exhibit unique structural elements compared to other EN classes, including the α7 and two disorder insertions indicated by dashed lines.

Structural comparison shows that KASV EN shares a similar overall architecture with ENs from other bunyaviruses and influenza virus (RMSD 2.7 to 3.5 Å, 42% to 63% residue coverage) (**Figure 3A and 3B**), despite their very low sequence homology (identity ranging from 11%-17%, similarity ranging from 16%-24%), suggesting conserved domain organization among viral cap-snatching endonucleases. Nevertheless, KASV and CCHFV ENs exhibit unique structural features compared to other viral ENs, including the α7 helix and one or two disorder insertions (insertions 1 and 2), which are exclusively presented in nairovirus ENs (**Figure 3B**). The α7 helix in both structures wraps the base of the central β-sheet, providing stable support to the overall structural module and partially sealing the bottom of the active site pocket. Insertion 1 locates between the β1 and β2 and is adjacent to the active site, which is relatively conserved and exists in nearly all nairoviruses. By contrast, insertion 2 follows α7 and connects to the outmost β3, positioned far away from the active site. This insertion is highly variable in length and sequence and is found only in CCHFV and other nairoviruses with a large L protein (**Figure S3C**). Together, these unique structural elements in nairoviruses ENs might play specific functional roles, such as regulating EN activity, facilitating substrate binding, or establishing interactions with other functional domains of viral polymerase.

### The active site of KASV EN exhibits a unique metal ion binding mode

The apo KASV EN structure shows no metal ions in the active site, whereas distinct electron density for two metal ions was observed in MnCl₂-soaked crystals. In the Mn^2+^-bound structure, the active site features a conserved E_668_…E_682_…P_718_D_719_…E_831_…K_841_ catalytic motif that binds two manganese ions (**Figure 4A**). Mn1 was octahedrally coordinated by the side chain of D719, the carbonyl group of V832, two discrete water molecules (W2 and W4), and two bridging water molecules (W1 and W3), which form hydrogen bonds with the side chains of E668 and E831. Similarly, Mn2 exhibits a predominantly solvent-mediated coordination sphere with D719 as its sole direct protein ligand. This coordination involves one discrete water molecule (W4), together with three bridging water molecules (W5, W6, and W7) interacting with the side chains of E831, E682, and E668. In addition, the putative catalytical residue K841 and the K848 associated with metal ion coordination via bridging water molecules (W3 and W8) (**Figure 4A**). Comparison of the apo and Mn^2+^-bound structures reveals subtle conformational changes in the active site residues upon metal ions binding, except for the E682 side chain in LoopE, which repositions toward the Mn2-binding site (**Figure S4A**). The apo structures of CCHFV and KASV ENs exhibit nearly identical active site conformations (**Figure S3B**). Sequence alignment analysis further reveals that the metal ion-coordination residues are strictly conserved among nairovirus ENs (**Figure S3C**), implying a common metal-ion binding configuration shared by nairovirus EN.

**Figure 4.**
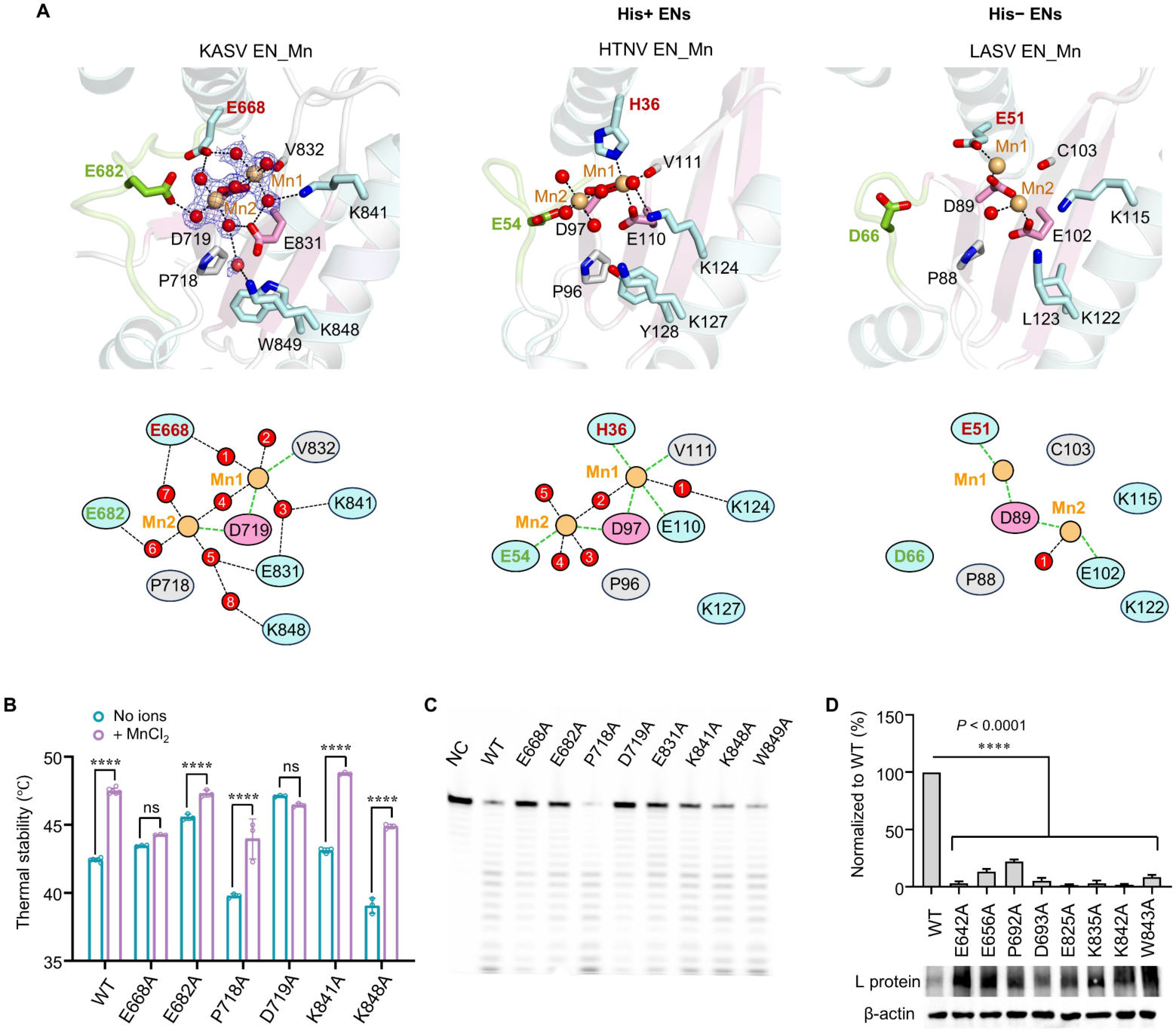
KASV EN features a unique two-metal-ion binding mode critical for catalytic activity. (A) Comparison of the active site and metal ion binding mode of KASV EN with those of His+ and His− endonucleases in the presence of manganese ions. Key residues involved in metal ion coordination are labeled and shown as sticks. Manganese ions and coordinated water molecules are shown as orange and red spheres, respectively, with coordination interactions indicated by black dashed lines. The *2F_o_*-*F_c_* electron density map (contoured at 0.7 σ) is overlaid for manganese ions and coordinated water molecules in KASV EN. The lower panel shows a schematic diagram of metal-ion binding mode in different ENs, with direct coordination shown as green dashed lines and indirect coordination as black dashed lines. PDB entries: 5IZE (HTNV), 5J1P (LASV). (B) Thermal stability of KASV EN WT and mutants with or without MnCl_2_. *T*_m_ values are represented as the mean values ± SD derived from three independent experiments. Statistical significance of differences between the no-ions and MnCl_2_ groups was evaluated via two-way ANOVA analysis. \*\*\*\**P* < 0.0001; ns, not significant. (C) Endonuclease activity of KASV EN WT and active site mutants. The 19-mer U-rich ssRNA was incubated with 1 μM WT or mutant protein in the presence of 2 mM MnCl_2_. As a negative control (NC), the ssRNA was incubated with WT protein in the absence of MnCl_2_. Some residual background activity may associate with sample contamination rather than KASV EN activity. (D) Transcriptional activity of WT and mutant CCHFV L proteins. BSR-T7/5 cells were transfected with T7 RNA polymerase-driven mini-replicon system along with two helper plasmids. eGFP reporter gene expression was quantified and normalized to that of WT group (set as 100%). The statistical significance was determined by two-tailed Student’ s *t* tests. \*\*\*\**P* < 0.0001. Data represent mean ± SEM of two independent experiments (n=2×3, with two biological replicates, each consisting of three technical replicates). Immunoblot analysis of 3×FLAG-tagged WT and mutant L proteins is shown. The β-actin was used as an internal control.

Comparison with previously reported Mn^2+^-bound His+ ENs and His− ENs structures reveals that KASV EN exhibits similar configurations for its active site residues. However, its metal ion binding mode is strikingly different compared to both enzyme classes (**Figure 4A**). First, KASV EN contains E668 (equivalent to His in His+ ENs and Glu/Asp in His− ENs) which simultaneously participates in Mn1 and Mn2 coordination via bridging waters, whereas its counterparts exclusively bind Mn1 directly (14,15,18,19). Second, LoopE in KASV EN maintains a closed conformation in both apo and Mn^2+^-bound structures, the Mn2 coordination is mediated solely by conformational adjustments of the side chain of E682 positioned above it (**Figure S4A**). Differently, the flexible loop exists in two distinct conformations in the other two enzyme classes: in His+ ENs, it undergoes an open-to-closed conformational transition upon metal ions or inhibitor binding(14,16), while in His− ENs, it adopts an open conformation incapable of metal coordination(18,20). Third, with the exception of D719, which directly coordinates with both Mn1 and Mn2, all other conserved residues in KASV EN indirectly coordinate with the metal ions via bridging water molecules. By contrast, in the other two enzyme classes, all these residues are nearly directly involved in metal ion binding. Collectively, these observations suggest a specific metal ion binding mode in the active site of KASV EN and imply a unique catalytic mechanism within nairovirus ENs.

### Two metal ions are required to achieve full catalytic activity of KASV EN

To corroborate our structural finding, we performed single-point alanine mutations at eight conserved active site residues to investigate their impacts on metal ion-induced protein stability and catalytic activity. As expected, mutating D719 completely abolished metal ion-binding stabilization (Δ*T*_m_ = −0.7 °C), consistent with its direct coordination with both Mn1 and Mn2. Mutations at E668 and E682, which indirectly participate in binding both metal ions, also significantly reduced the stabilization effect (Δ*T*_m_ = 0.8, and 1.7 °C, respectively), indicating that these residues are essential for optimal metal binding. By contrast, mutations at P718, K841 and K848 had a minimal impact on Mn^2+^-induced protein stability (**Figure 4B and Table 1**), likely due to their limited contribution to metal ion binding. The E831A and W849A mutants displayed noncanonical thermal denaturation curves (**Figure S5B)**, presumably caused by protein aggregation or conformational instability during thermal ramping.

**Table 1.** Thermal stability, Mn^2+^ -induced stability, endonuclease activity of KASV EN mutants, and transcriptional activity of CCHFV L mutants.

| Constructs | Thermal stability (°C) <sup>a</sup> |  |  | Endonuclease activity <sup>b</sup> | Transcriptional activity <sup>c</sup> |
| --- | --- | --- | --- | --- | --- |
| | No ion | Mn<sup>2+</sup> | $\Delta T_m$ | | |
| WT | 42.4 ± 0.1 | 47.5 ± 0.2 | 5.1 | 100 | 100 ± 2 |
| E668A | 43.5 ± 0.1 | 44.3 ± 0.0 | 0.8 | 36 | 3 ± 2 |
| E682A | 45.6 ± 0.2 | 47.3 ± 0.3 | 1.7 | 57 | 13 ± 3 |
| P718A | 39.8 ± 0.2 | 43.9 ± 1.5 | 4.1 | 124 | 22 ± 2 |
| D719A | 47.1 ± 0.1 | 46.4 ± 0.1 | -0.7 | 30 | 5 ± 3 |
| E831A | NA | NA | NA | 56 | 2 ± 1 |
| K841A | 43.1 ± 0.2 | 48.7 ± 0.1 | 5.6 | 77 | 3 ± 2 |
| K848A | 39.0 ± 0.5 | 44.9 ± 0.2 | 5.9 | 95 | 2 ± 1 |
| W849A | NA | NA | NA | 110 | 8 ± 2 |

<sup>a</sup> $T_m$ values calculated from DSF assay in the absence of ions, the presence of 2 mM MnCl<sub>2</sub>.
<sup>b</sup> Measured by *in vitro* endonuclease activity assay. Activity was determined by quantifying the percentage of substrate ssRNA degraded. Values for active site mutants are normalized to the activity of the WT (set as 100%).
<sup>c</sup> Measured by CCHFV mini-replicon assay. Activity is normalized to the WT and presented as percentages.
NA, not applicable.

In parallel, we investigated the endonuclease activity of these mutants using the 19-mer U-rich ssRNA substrate in the presence of Mn^2+^. Mutation in the key metal ion-coordination residues (E668, E682, D719, and E831) dramatically reduced EN activity. While the putative catalytic residue K841A retained activity, albeit at a lower level than the WT, suggesting that it contributes to but is not essential for EN activity. Mutations at P718, K848, and W849 had no effect on EN activity (**Figure 4C and Table 1**), in line with their minimal or no involvement in metal ion coordination. Taken together, these findings indicate that the conserved residues involved in coordinating the two metal ions are critical for both metal ion binding and EN catalytic activity.

To get deeper structural insights into the relationship between metal ion binding and EN activity, we crystallized three key mutants (E668A, E682A, and D719A,) and successfully determined their high-resolution structures. While the D719A mutant completely abolished metal ion binding in the active site, both E668A and E682A mutants retained binding to only a single metal ion, corresponding to Mn1(**Figure S4B**). These structural observations are fully consistent with the biochemical data from thermal stability and EN activity assays. Together, the combined biochemical and structural data highlights D719 as a critical residue for coordinating both metal ions and for catalytic activity, supporting the notion that binding of both metal ions is essential for the full enzymatic function.

### The essential role of the KASV EN active site in viral transcription

To assess the functional relevance of KASV EN in viral transcription, we further investigated the roles of the active site residues in the context of full-length RNA polymerase. Since no mini-replicon system is available for KASV, we used an established CCHFV mini-replicon system in which a GFP reporter gene serves as a readout of the transcription activity of the viral polymerase (43). The WT L protein exhibited robust transcription activity. However, mutating the metal ion-coordination residues E642, E656, D693, E825, and K835 (homologous to KASV EN E668, E682, D719, E831, and K841) nearly abrogated the transcription activity, consistent with their impaired EN activity *in vitro*. Interestingly, mutations at the residues P692, K842, and W843 (homologous to KASV EN P718, K848, and W849) also caused a significant reduction in activity (**Figure 4D**); although these residues are minimally implicated in metal ion binding and do not affect EN activity *in vitro*, implying their additional functional roles in transcription beyond EN activity. Collectively, these data suggest that transcription by the full-length RNA polymerase is highly dependent on the two-metal ion-dependent endonuclease activity.

### Inhibition of KASV and CCHFV endonuclease activity by DPBA, L-742,001 and BXA

As described above, three representative inhibitors (DPBA, L-742,001, and BXA) (**Figure 5A**), were shown to enhance the stability of KASV EN. To further evaluate their binding affinity and inhibitory potential against nairovirus ENs, we conducted microscale thermophoresis (MST) assays and endonuclease activity experiments. KASV EN displayed the highest affinity to BXA, with a dissociation constant of Kd = 4.5 μM, which is about fifteen and thirty times lower than those of DPBA (Kd = 70.7 μM) and L-742,001 (Kd = 140.1 μM), respectively (**Figure 5B**). In the EN activity assays, both L-742,001 and BXA efficiently inhibited KASV EN activity at concentrations of 400 and 200 μM, respectively, whereas DPBA showed no detectable inhibition at 400 μM under the tested conditions (**Figure S6B**). Dose-dependent assays revealed that L-742,001 had a half-maximal inhibitory concentration (IC_50_) of 176.5 μM for KASV EN, approximately 10 to 100-fold higher than those reported for other bunyavirus ENs (17,31,44). BXA exhibited pronounced higher inhibitory activity than L-742,001, with an IC_50_ of 28 μM (**Figure 5C**). Nonetheless, its potency remains significantly lower than the nanomolar-range IC_50_ reported for influenza virus ENs (26). Parallel assays with CCHFV EN produced similar binding and inhibition profiles to those observed for KASV EN (**Figure S6**). The MST and enzymatic activity data confirm that both L-742,001 and BXA can bind to and inhibit nairovirus ENs *in vitro*, with BXA exhibiting superior binding affinity and inhibitory efficacy.

**Figure 5.**
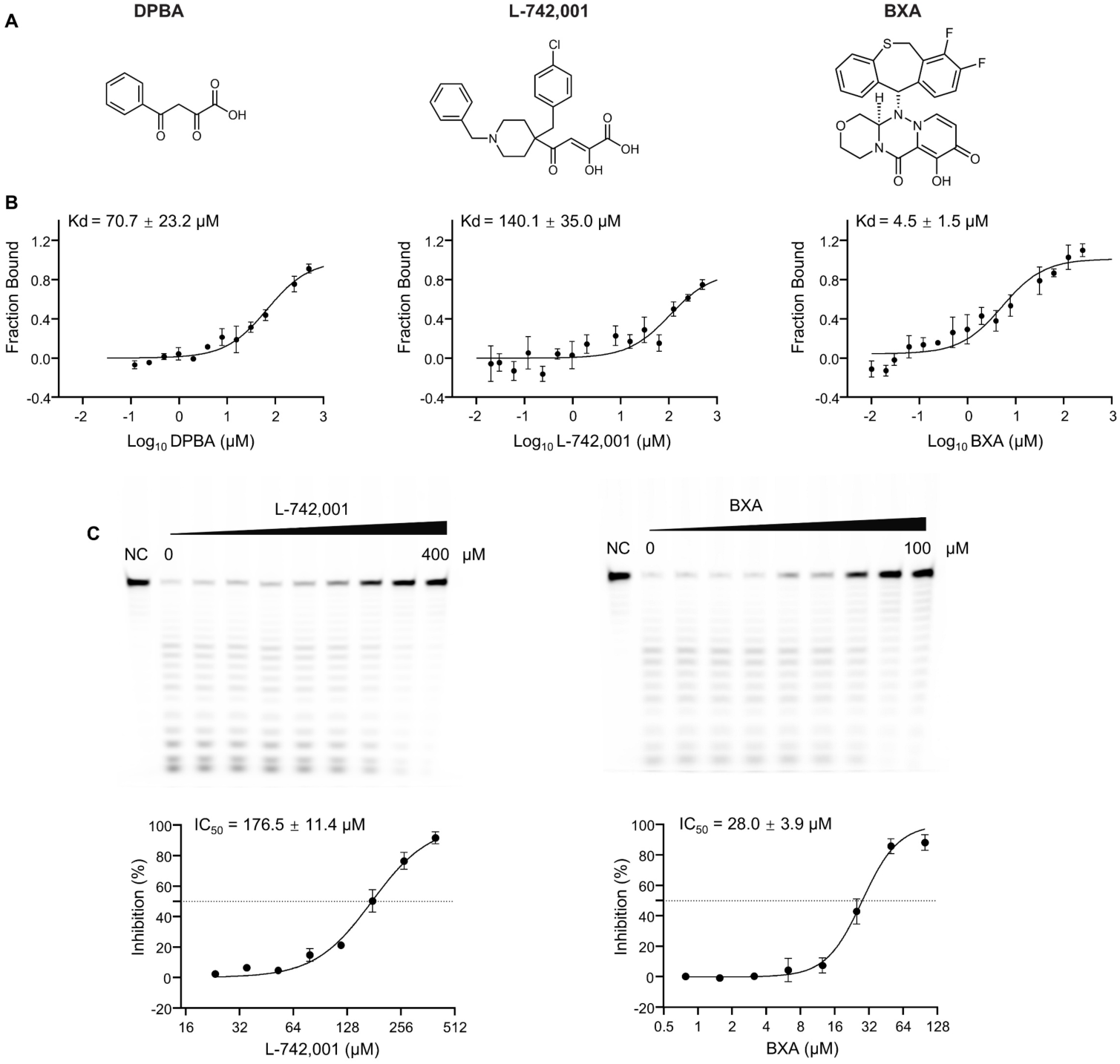
Binding affinity and inhibitory effect of three representative inhibitors on KASV EN. (A) Chemical structures of DBPA, L-742,001, and BXA. (B) MST binding assays of KASV EN with DBPA, L-742,001, and BXA. Dissociation constants (Kd ± SD) were calculated from three independent experiments. Note that the data point in L-742,001 MST curves do not reach a maximum plateau, as the high centration L-742,001 (1 mM) caused precipitation of KASV and CCHFV ENs (also see Figure S6A). (C) Dose-dependent inhibition of KASV EN activity by L-742,001 and BXA. A 1.5 μM of KASV EN was incubated with 19-mer U-rich ssRNA at 37°C in the presence of 2 mM MnCl_2_ and increasing concentrations of each inhibitor. Experiments were performed in triplicate, and a representative gel image is shown. Data represents the mean ± SD from three independent experiments.

### Binding modes of the inhibitors in KASV EN

To elucidate the mechanisms of action of DPBA, L-742,001 and BXA against nairovirus ENs, we co-crystallized KASV EN with each inhibitor. The three co-crystal structures were determined at resolutions of 2.15 Å, 1.95 Å, and 2.16 Å, respectively. All three structures displayed unambiguous electron density for the compounds, which chelate the two manganese ions at the active site center via their respective head groups (diketo acid in DPBA, L-742,001, and an oxazino-pyridotriazin-dione moiety in BXA) in a similar fashion (**Figure 6A**). In each complex, Mn1 and Mn2 are coordinated by the side chain of the central D719, two adjacent and planar oxygen from the inhibitor’s head group, and bridging water molecules that interact with the conserved active site residues. Beyond the metal chelating moiety, the remaining portions of the inhibitors adopt distinct conformations. The phenyl group of DPBA makes no direct contact with the protein, and only weak electron density was observed for this group. In the L-742,001-complexed structure, the piperidine central ring is oriented perpendicular to the diketo acid group with well-defined electron density, while the phenyl and *p*-chlorobenzene moieties appear disordered, possibly due to a lack of interactions with the surrounding residues. Notably, the BXA-bound structure exhibits well-resolved electron density across the entire molecule. In addition to coordinating the manganese ions, BXA’s V-shaped tail group engages in additional interactions with residues distal to the catalytic center (**Figure 6A**). Specifically, R661 packs against the V-shaped tail group, forming a cation-π stacking with the aromatic ring of BXA, while L660 and G664 contribute hydrophobic contacts. These interacting residues are highly conserved among nairovirus ENs, highlighting their importance for further inhibitor exploration (**Figure S3C**). Together, these structural findings demonstrate that BXA establishes more extensive and stable interactions with KASV EN than DPBA and L-742,001, accounting for its superior binding affinity and inhibitory potency.

**Figure 6.**
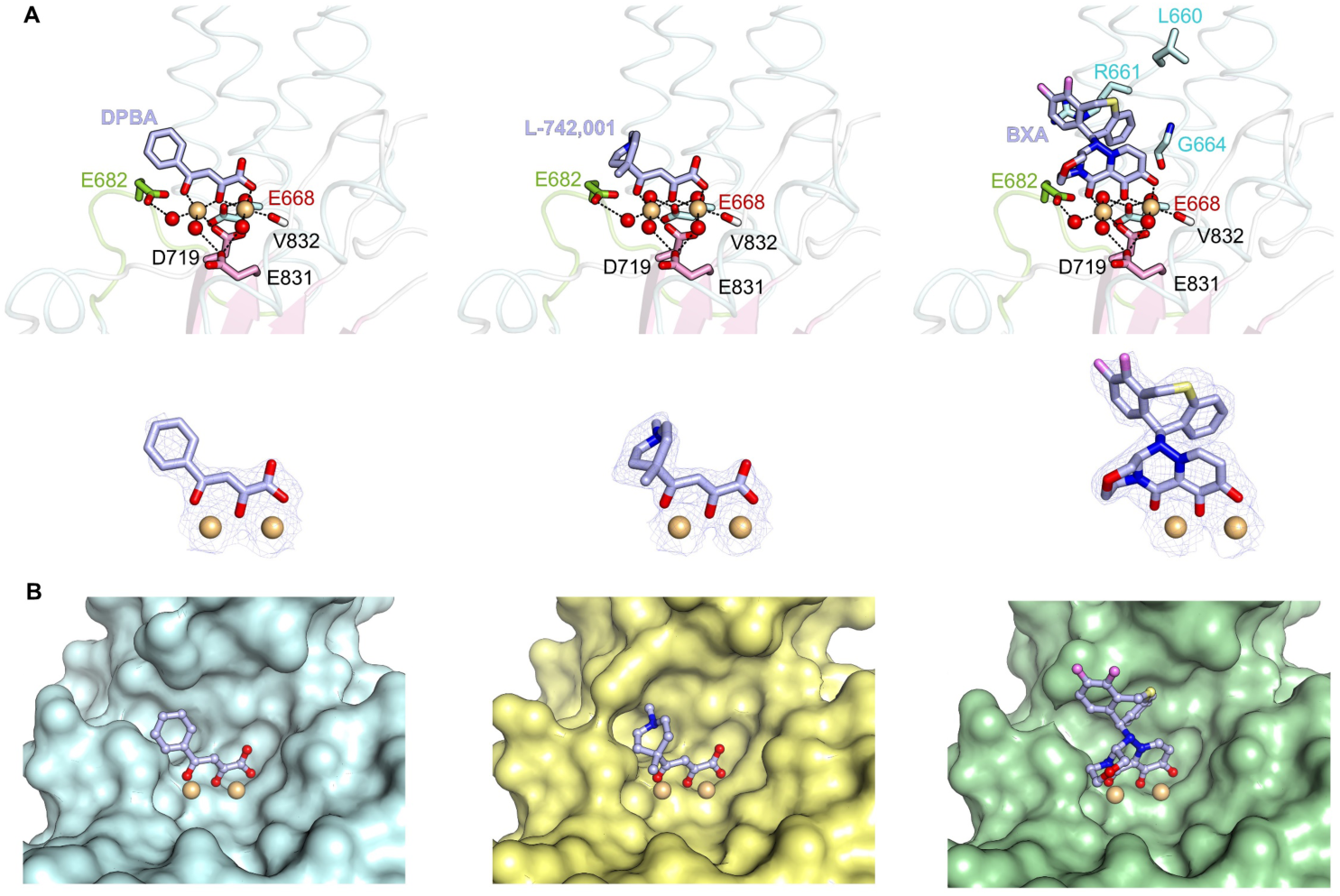
Crystal structures of KASV EN in complex with DBPA, L-742,001, and BXA. (A) Detailed interactions between KASV EN and the inhibitors. The inhibitors are shown as purple sticks, with *2F_o_*-*F_c_* electron density maps (contoured at 0.8 *σ)* displayed in the lower panel. Key residues involved in inhibitor binding are labeled and shown as sticks. Bound manganese ions and water molecules are shown as orange and red spheres, respectively. BXA forms additional interactions with the pocket-lining residues, which are highlighted in cyan. (B) Surface representation of the inhibitor-binding pocket in the KASV EN.

## Discussion

The cap-snatching endonucleases encoded by sNSVs belong to the PD-(D/E)xK nuclease superfamily that requires divalent metal ions for catalysis. Unlike other viral endonucleases that situated at the N terminus of the L protein, the nairovirus EN domain was previously predicted to be located around amino acids 700 in the L protein (20), and its function was first identified using a CCHFV virus-like particle system, in which the active-site signature residue D693 was shown to be necessary for mRNA transcription (45). A subsequent study showed that the CCHFV and other related nairovirus ENs are enzymatically inactive *in vitro*, despite being capable of binding metal ions via conserved acidic residues (E642 and E656) (22). In our study, we found the isolated CCHFV and KASV ENs only exhibited low *in vitro* activity toward 19-mer U-rich ssRNA, exclusively in the presence of Mn^2+^, this appears to slightly discrepancy with the previous results, which might be due to the different experimental conditions. Nevertheless, the isolated nairovirus ENs show minimal or low activity *in vitro*, suggesting that efficient EN activity likely requires the structural or functional context of the full-length RNA polymerase.

Using the antibody-assisted crystallization strategy, we clearly depicted both the apo and metal ion-bound states of nairovirus endonuclease for the first time by determining high-resolution structures of KASV and CCHFV EN. Unlike other known viral ENs, KASV EN exhibits a unique two-metal-ion binding mode (**Figure 4A**), in which the conserved D719 serves as the sole direct ligand coordinating both metal ions, while the remaining active site residues bind the metal ions indirectly via bridging water molecules. This particular metal ion coordination mode likely arises from the native active-site configuration, which is dictated by the spatial arrangement of the conserved residues. Notably, conservation analysis reveals that the active site architecture is highly conserved among nairovirus ENs, composed of residues that are evolutionarily preserved (**Figure S7**), strongly suggesting a shared metal ion binding mechanism within the *Nairoviridae* family. Our data also provide structural evidence that nairovirus ENs likely represent a distinct class of cap-snatching endonucleases, aligning with earlier proposals that they take a unique evolutionary position between His+ and His− endonucleases(22). For sNSV-encoded endonucleases, the two-metal-ion catalysis mechanism was initially proposed for influenza virus EN and supported by our structural and enzymatic studies on the Ebinur Lake virus EN(15,19). In this study, mutations at residues directly or indirectly involved in both manganese ions coordination significantly impaired the KASV EN activity (**Figure 4C**), and the replication and transcription activity in a CCHFV based mini-replicon system (**Figure 4D**). Structural analyses of the inactive mutants show either partial or complete loss of metal ion binding (**Figure S4B**). These findings indicate that EN activity is highly dependent on the binding of two metal ions, suggesting nairovirues ENs employ a two-metal-ion catalysis akin to other cap-snatching endonucleases, albeit through distinct metal ion coordination mode.

The two-metal-ion catalysis has been established as a general mechanism for many nucleases, with both metal ions (referred to as metal A and B) playing distinct roles in the catalytic cycle (46–50). On one hand, metal A can activate the hydroxyl nucleophile to facilitate nucleophile attack on the scissile phosphodiester of the nucleic acid substrates. Water molecules, bound to metal A, act as the most common nucleophiles and can be directly activated by the metal ion or deprotonated by a general base. On the other hand, both metal A and metal B stabilize the ground-state, transition-state, and post-reactive-state by interacting with the bridging or nonbridging oxygen atoms of the scissile phosphate. Remarkably, the two metal ions can reposition relative to each other during different reaction stages, adopting an optimal spatial coordination geometry that favors the nucleophile attack and product formation. The realization of the above-mentioned functions of the metal ions involves the dissociation of metal ions from the original ligands and the re-coordination with new ligands. Based on this, we postulate two functional roles of water molecules in the metal ion binding mode of KASV EN. First, one metal ion-bound water might serve as a nucleophile, especially the water bound with Mn1, which occupies a similar spatial position as metal A (**Figure S8**). Second, the bridging water molecules between Mn^2+^ and binding residues may reduce the free energy barrier of ligand binding by acting as a lubricant, thereby accelerating rapid ligand exchange to achieve efficient catalysis.

Cap-snatching endonucleases have emerged as promising targets for antiviral drug development. We systematically evaluated the efficacy of three representative inhibitors against KASV and CCHFV ENs. Our results show that BXA exhibits the highest binding affinity and the most potent inhibitory activity among the three tested compounds (**Figure 5 and Figure S6**). Structural analyses of EN-inhibitor complexes further reveal that all three inhibitors chelate the two catalytic metal ions in a similar manner. However, BXA uniquely forms additional stabilizing interactions with the enzyme (**Figure 6**), which likely contribute to its superior inhibitory efficacy. Despite its effectiveness, BXA inhibits both KASV and CCHFV ENs with micromolar IC_50_ values, markedly less potent than its nanomolar-level inhibition of influenza virus EN. By comparing the binding mode of BXA in IAV and KASV EN, we observed that the IAV EN features a relatively compact and narrow active site pocket, enabling BXA to nearly fully occupy the pocket and form extensive interaction networks with surrounding residues (**Figure S9**). By contrast, the active site pocket of KASV and CCHFV ENs is relatively large and quite open, resulting in limited residue contacts and only partial occupancy by BXA (**Figure 6B**). This structural distinction in the binding pocket likely explains BXA’s higher efficacy against IAV. Notably, the EN of nairovirus and other bunyavirus families form a common open active site pocket, posing a challenge for inhibitor design. Nevertheless, a previous study has demonstrated that BXA can effectively inhibit CCHFV infection in cells with a IC_50_ at the micromolar level, and its sodium form significantly improves survival rates in a mouse model(43), highlighting its potential as a drug candidate for anti-CCHFV. Given the spatial vacancy within the BXA binding pocket, future optimization efforts should focus on modifying the BXA scaffold to extend into adjacent sub-pockets and establish additional stabilizing interactions with the pocket-lining residues.

In summary, our work offers the first atomic-resolution insight into nairovirus ENs. A series of high-resolution structures of CCHFV and KASV ENs reveal a distinctive two-metal ion binding mode critical for enzymatic function. This unique coordination involves multiple bridging water molecules with specific implications for efficient catalysis and adaptation to diverse substrates or environmental conditions. Importantly, this two-metal ion mode is conserved across nairovirus ENs and is susceptible to inhibition by existing endonuclease-targeting compounds. These findings provide a critical foundation for developing novel, potent, and broad-spectrum antivirals targeting CCHFV and other related nairoviruses.

## Data availability

Atomic coordinates and structure factors for the reported crystal structures have been deposited with the Protein Data bank under accession numbers 9UZA, 9UZB, 9UZC, 9UZD, 9UZE, 9UZF, 9UZG, 9UZH, and 9UZI.

## Acknowledgments

We thank Dr. Peng Gong for crystallography experimental equipment and discussions. We thank the Center for Instrumental Analysis and Metrology of Wuhan Institute of Virology for providing technical assistance. We thank the staff of the BL02U1 and BL10U2 beamlines at Shanghai Synchrotron Radiation Facility (Shanghai, People’s Republic of China) for assistance during X-ray diffraction data collection. This work was supported by National Key Research and Development Program of China (2022YFC2303300) and the CAS Pioneer Hundred Talents Program (E2YC0001) to Z.D., the National Natural Science Foundation of China (U22A20336) to Z.H., the National Natural Science Foundation of China (32400129), Hubei Province Natural Science Foundation (2022CFB881), and Natural Science Foundation of Wuhan (2024040801020249) to W.K.

## Author contributions

Z.D., W.K., H.Z., and M.W. conceived and supervised the project; W.K. performed biochemical and crystallization experiments with the help of Z.T., J.L., and X.Z.; Z.T. performed MST assays; W.K., Z.D. and Z.T. determined the crystal structures; G.Z. conducted mAb generation experiments; F.W. performed mini-replicon assay; J.T. conducted western blotting experiments; H.Zhang performed protein topology and phylogenetic analyses; W.K. and Z.D. analyzed the data and prepared the manuscript with input from all authors. Declaration of interests: The authors declare that they have no competing interests.

