## Supplemental figures and tables for "Structure and function of the nairovirus cap-snatching endonuclease"

### Supplemental information

### Figures S1 to S9, and Table S1

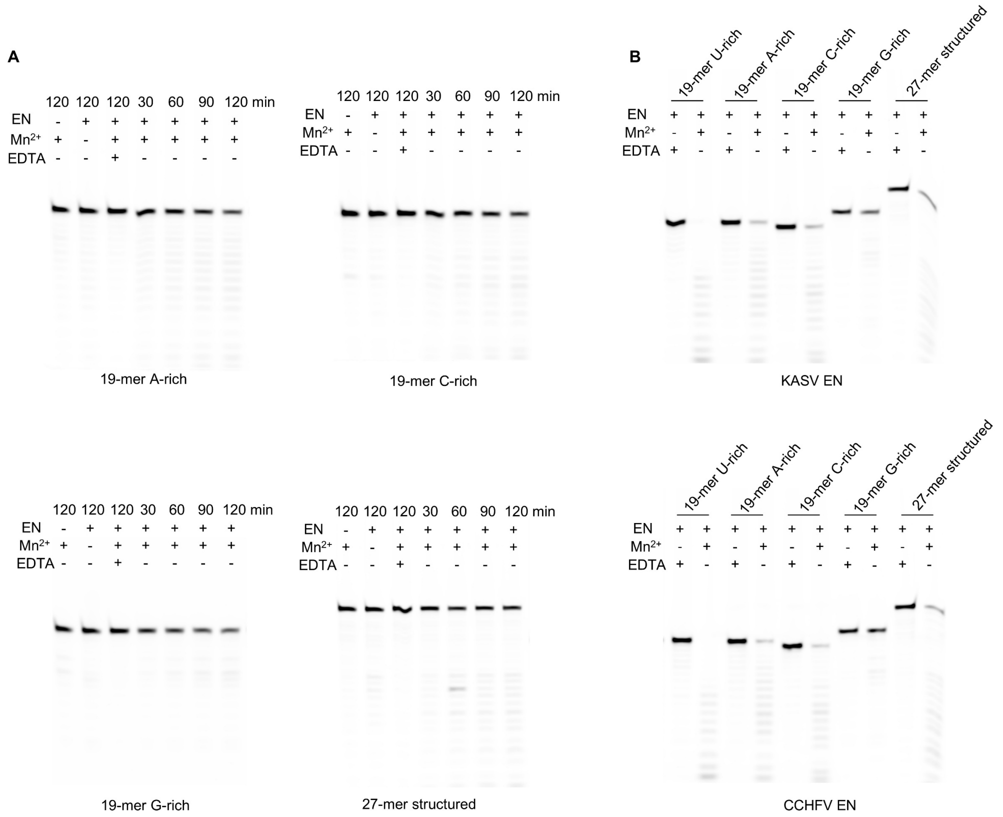


**Figure S1. RNA substrate preference of KASV and CCHFV ENs.** (A) Time course of endonuclease activity of KASV EN with different ssRNA substrates. All assays were performed with 1 μM enzyme in the presence of 2 mM $MnCl_2$. (B) Endonuclease activity of KASV and CCHFV EN at high enzyme concentrations. Reactions were conducted with KASV EN at 2 μM or CCHFV EN at 4 μM in the presence of 2 mM $MnCl_2$ and various ssRNA substrates.

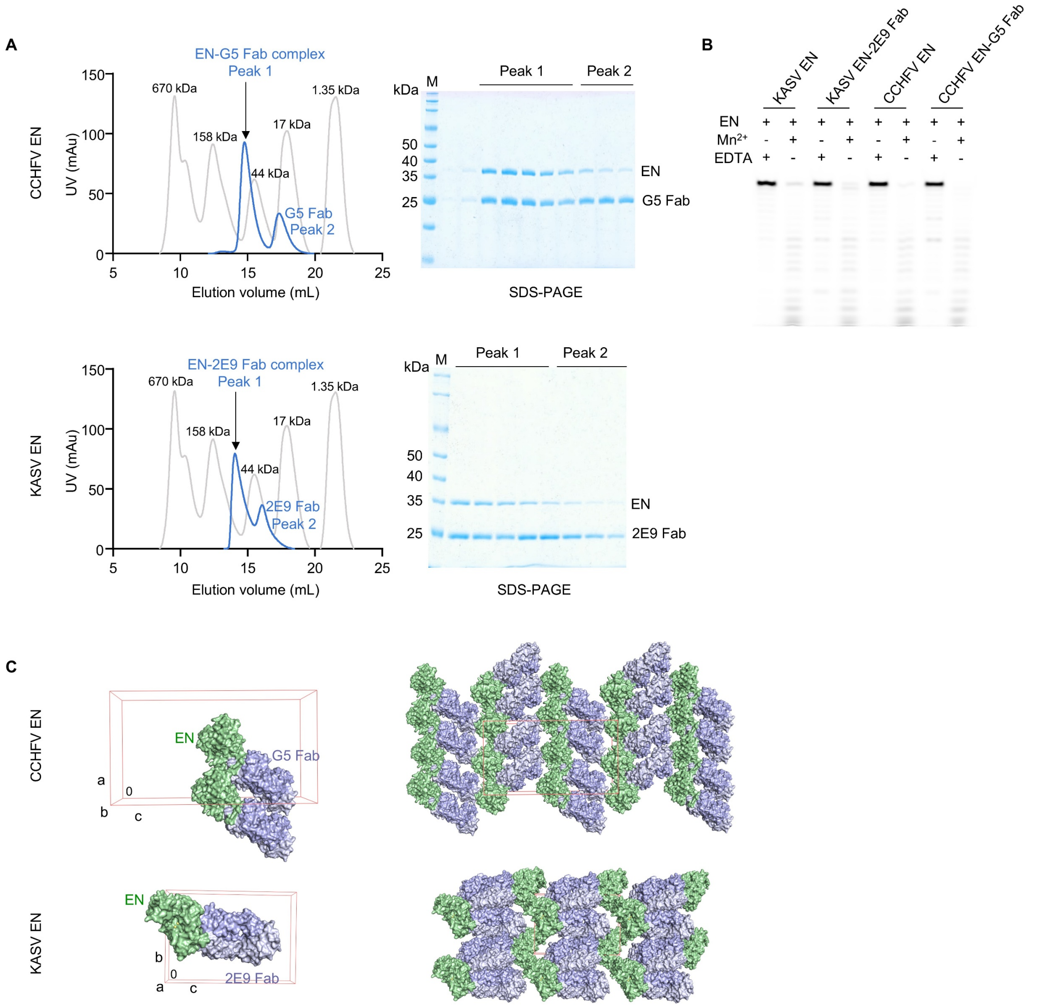


**Figure S2. Antibody-assisted crystallization of CCHFV and KASV ENs.** (A) Size-exclusion (Superdex 200 Increase 10/300 GL) and SDS-PAGE profiles of the purified EN-Fab complex. The gel filtration profiles of EN-Fab complex and standard proteins are colored in blue and grey, respectively. (B) The G5 and 2E9 Fab has no effects on the endonuclease activity of CCHFV and KASV ENs. Reactions were conducted with protein at 1 μM (KASV EN and EN-2E9 Fab complex) or 2 μM (CCHFV EN and EN-G5 Fab complex) in the presence of 2 mM $MnCl_2$ and 19-mer U-rich ssRNA ssRNA substrates. (C) Antibody-mediated crystal packing in CCHFV and KASV ENs crystals. The unit cell is shown in a red cube, with EN-Fab complex molecules in the crystallographic asymmetric unit represented as surfaces.

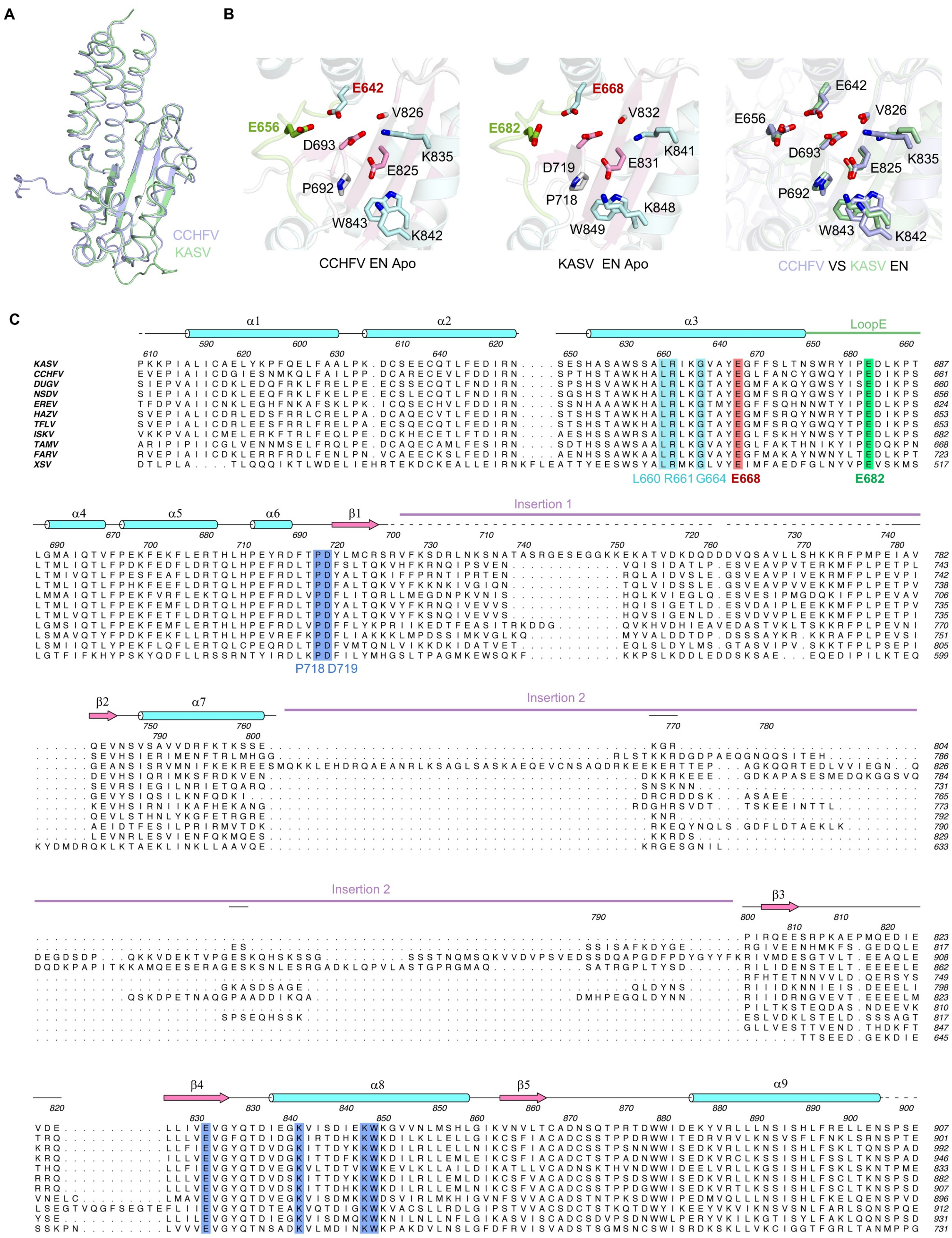


**Figure S3. CCHFV EN shares a highly similar overall fold and active site configuration with KASV EN.** (A) Superposition of overall structure of the apo form of CCHFV and KASV ENs. (B) Comparison of the active site of CCHFV and KASV ENs. Coloring scheme is the same as in Figure 4A. (C) Sequence alignment of representative nairovirus ENs. The key conserved active site residues involved in manganese ions coordination and BXA binding are highlighted in different colors. The two insertions are indicated with purple straight lines. The secondary structure of KASV EN is shown over the sequence alignment, with α-helices and β-strands shown as springs and arrows, respectively. The amino acid numers is shown, with the upper and lower rows corresponding to the CCHFV and KASV EN, respectively.HAZV, Hazara virus; TFLV, Tofla virus; ISKV, Issyk-Kul virus; TAMV, Tamdy virus; FARV, Farallon virus; XSV, Xinzhou Spider Virus.

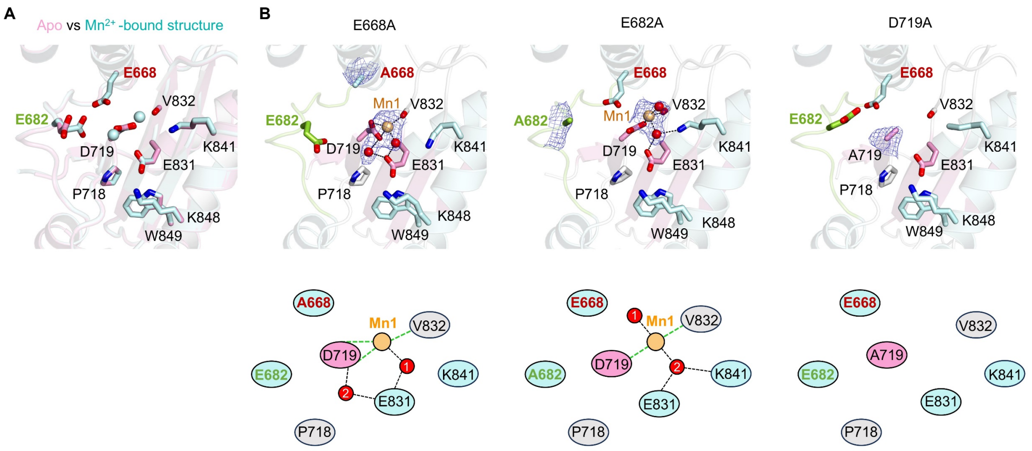


**Figure S4. Comparison of the active site of the apo form, $Mn^{2+}$-bound form, and mutants of KASV EN.** (A) Superposition of the active site of apo and $Mn^{2+}$-bound KASV EN structures. (B) The active site architecture of three KASV EN mutants. $2F_o$-$F_c$ electron density maps (contoured at 0.7 σ) are overlaid for the mutated residues, bound manganese ions, and coordinated water molecules. Coloring scheme is the same as in Figure 4A. Schematic diagrams of metal ion binding mode for the mutants are shown in the lower panel.

Fig.S5

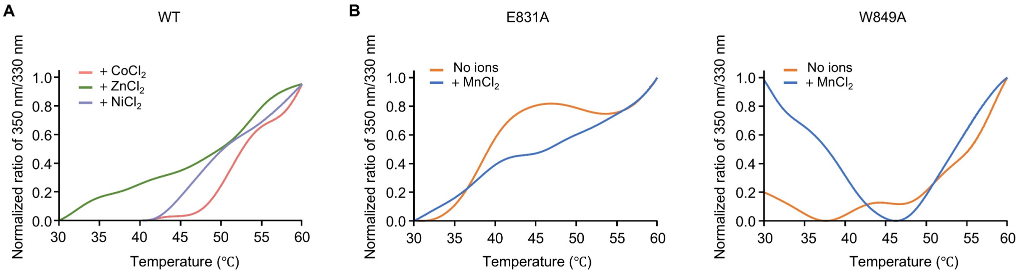


**Figure S5. Thermal denaturation profiles for KASV EN in the presence of metal ions ($Co^{2+}$, $Zn^{2+}$ or $Ni^{2+}$) (A), and for the E831A and W849A mutants (B).** The denaturation curves are noncanonical, which might be attributed to protein aggregation or conformational instability.

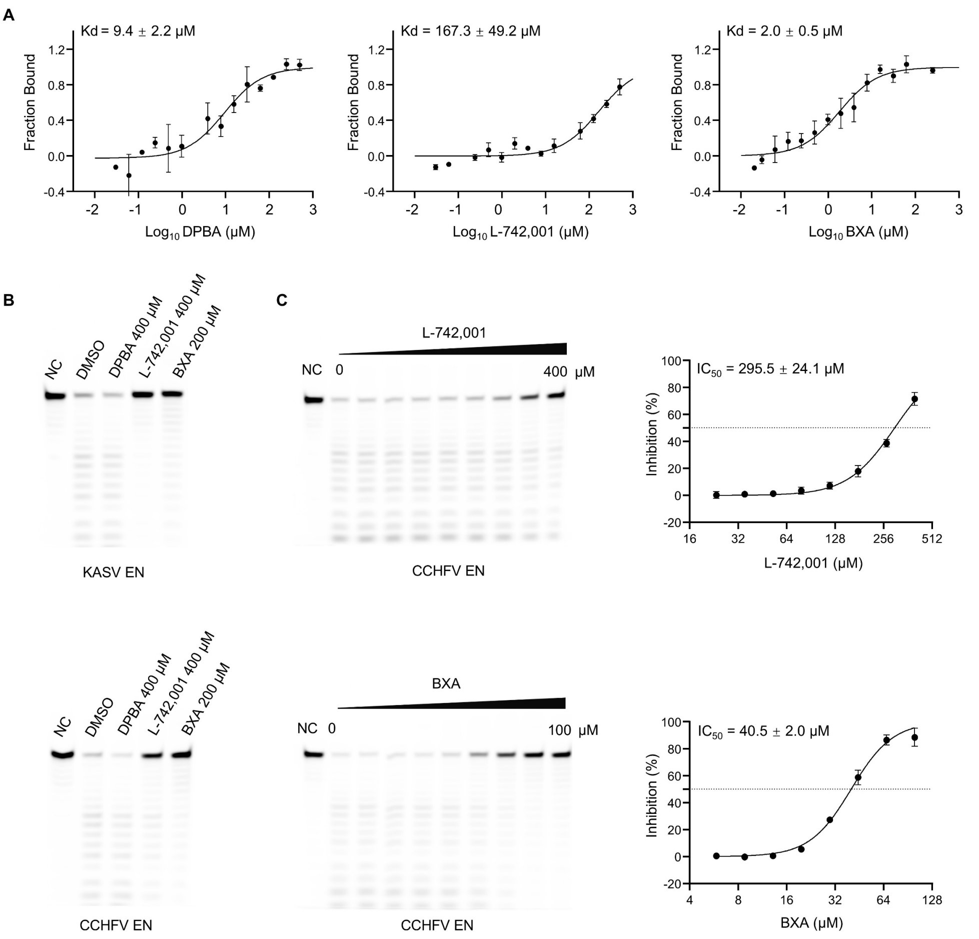


**Figure S6. Binding affinity and inhibition of three representative inhibitors against CCHFV EN.** (A) MST assays show the binding affinity between CCHFV EN and inhibitors (DPBA, L-742,001, and BXA). Data are presented as the mean value ± SD of three independent experiments. (B) Inhibition of KASV and CCHFN EN activity by DPBA, L-742,001, and BXA at higher concentration. The enzyme (KASV EN at 1.5 μM, and CCHFV EN at 3 μM) was incubated with 19-mer U-rich ssRNA at 37°C in the presence of 2 mM $MnCl_2$ and different inhibitors at indicated concentration. The reactions without $MnCl_2$ and with 5% vol/vol DMSO were set as the negative and positive controls, respectively. (C) Dose-dependent inhibition of CCHFV EN activity by L-742,001 and BXA. The experiments were performed as described for figure S6B, conducted in triplicate, and a representative gel image is presented. Data are shown as the mean ± SD from three independent experiments.

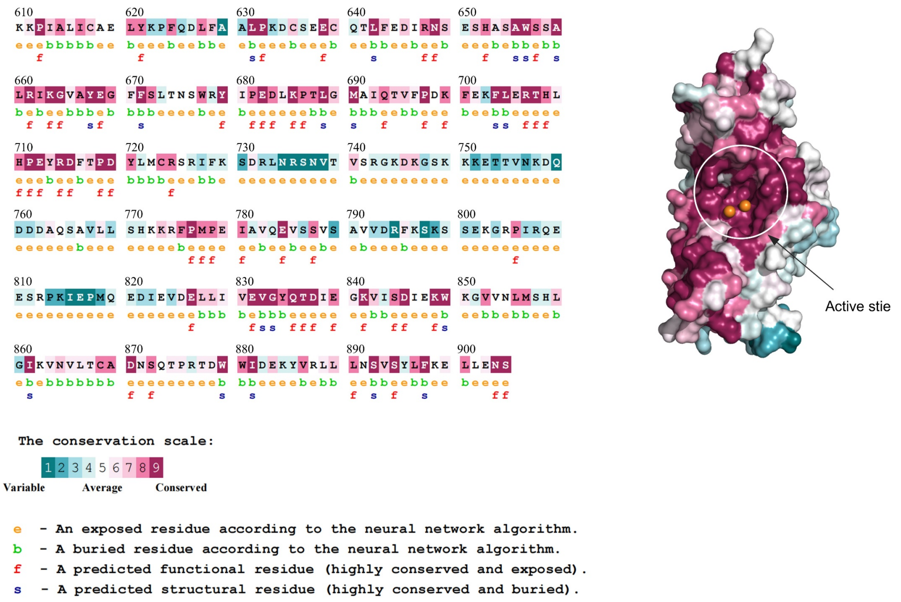


**Figure S7. Evolutionary conservation analysis of nairovirus ENs.** A multiple sequence alignment of 52 EN proteins from the *Nairoviridae* family shows that the active site architecture is highly conserved. Residues are colored-coded based on their conservation level from turquoise for variable to maroon for highly conserved. The conservation scores were calculated using an empirical Bayesian methodology by the ConSurf Server (53). The UniProt accession numbers and sequence boundaries of the 52 nairovirus ENs are listed in Table S3.

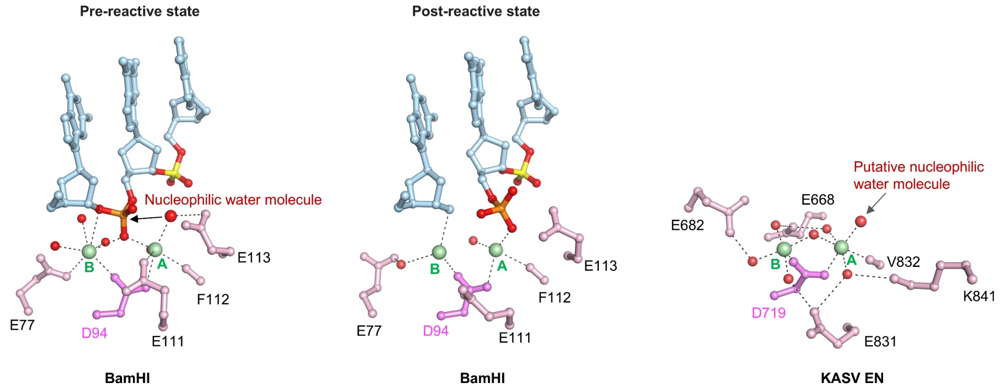


**Figure S8. Putative role of the metal ion-bound water molecules in the KASV EN catalysis.** The pre- and post-reactive states of BamHI (a type II restriction endonuclease) support a two-metal-ion catalytic mechanism. Structural analysis shows that a metal A (Mn1)-bound water molecule in the KASV EN active site might serve as a putative nucleophilic water molecule for catalysis. The conserved metal ion-coordination residues are shown as pink sticks, with the strictly conserved aspartate (D) highlighted in violet. The two metal ions are shown in green spheres and labeled as “A” and “B”. For clarity, the scissile phosphate is shown in orange, and the nucleophilic water molecule is represented as an enlarged red sphere. PDB entries: 2BAM and 3BAM.

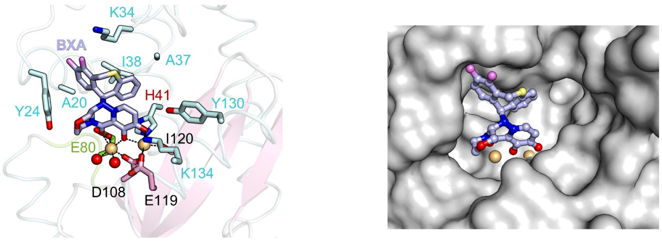


**Figure S9. Binding mode of BXA in IAV EN.** Detailed interactions (left panel) and binding pocket (right panel) of BXA with IAV EN are shown. Coloring scheme is the same as in Figure 6A for comparison. Compared to KASV EN, IAV EN features a more closed and narrower active site pocket that better accommodates BXA binding. PDB entry: 6FS6.

### Table S1. X-ray diffraction data collection and structure refinement statistics

| PDB code | CCHFV EN-Apo (9UZA) | KASV EN-Apo (9UZB) | KASV EN-$Mn^{2+}$ (9UZC) | KASV EN E668A (9UZD) | KASV EN E682A (9UZE) | KASV EN D719A (9UZF) | KASV EN-DPBA (9UZG) | KASV EN-L-742,001 (9UZH) | KASV EN-BXA (9UZI) |
|---|---|---|---|---|---|---|---|---|---|
| **Data collection** | | | | | | | | | |
| Space group | $P2_12_12_1$ | $P2_1$ | $P2_1$ | $P2_1$ | $P2_1$ | $P2_1$ | $P2_1$ | $P2_1$ | $P2_1$ |
| Cell dimensions | | | | | | | | | |
| a, b, c (Å) | 98.7, 99.8, 180.4 | 37.2, 90.0, 115.2 | 37.2, 80.7, 115.0 | 37.2, 80.3, 116.3 | 37.2, 81.1, 115.9 | 37.2, 81.2, 116.8 | 37.2, 80.0, 115.8 | 37.4, 80.3, 116.2 | 37.5, 80.1, 115.8 |
| α, β, γ (°) | 90.0, 90.0, 90.0 | 90.0, 93.5, 90 | 90.0, 93.5, 90.0 | 90.0, 97.8, 90.0 | 90.0, 93.4, 90.0 | 90.0, 97.7, 90.0 | 90.0, 93.2, 90.0 | 90.0, 93.4, 90.0 | 90.0, 93.4, 90.0 |
| Resolution (Å) [a] | 90.21–3.05 (3.21–3.05) | 28.41–1.90 (1.94–1.90) | 29.96–1.98 (2.02–1.98) | 29.30–2.40 (2.49–2.40) | 29.85–2.41 (2.50–2.41) | 29.33–2.47 (2.57–2.47) | 29.78–2.15 (2.22–2.15) | 29.92–1.95 (2.00–1.95) | 29.97–2.16 (2.23–2.16) |
| $R_{merge}$ | 0.149 (3.537) | 0.141 (0.451) | 0.118 (0.327) | 0.087 (0.516) | 0.071 (0.399) | 0.105 (0.658) | 0.105 (0.353) | 0.070 (0.479) | 0.098 (0.498) |
| $R_{meas}$ | 0.161 (3.819) | 0.192 (0.613) | 0.167 (0.462) | 0.111 (0.672) | 0.100 (0.561) | 0.139 (0.873) | 0.148 (0.495) | 0.098 (0.666) | 0.139 (0.704) |
| $CC_{1/2}$ | 0.998 (0.434) | 0.972 (0.629) | 0.973 (0.874) | 0.991 (0.652) | 0.994 (0.675) | 0.988 (0.678) | 0.984 (0.803) | 0.994 (0.606) | 0.987 (0.713) |
| $I/\sigma I$ | 5.1 (0.9) | 6.5 (2.5) | 5.0 (2.6) | 9.6 (2.1) | 10.8 (2.3) | 7.5 (1.7) | 4.8 (2.6) | 8.2 (2.6) | 5.3 (2.2) |
| Completeness (%) | 100.0 (100.0) | 99.5 (98.3) | 97.3 (89.0) | 94.9 (79.1) | 92.4 (78.7) | 99.6 (99.9) | 90.2 (99.0) | 98.3 (98.6) | 89.4 (97) |
| Redundancy | 13.1 (13.6) | 3.3 (3.3) | 2.8 (2.8) | 4.2 (3.0) | 3.1 (2.3) | 3.9 (3.9) | 3.0 (3.0) | 2.8 (2.8) | 2.6 (2.5) |
| **Refinement** | | | | | | | | | |
| Resolution (Å) | 3.05 | 1.90 | 1.98 | 2.40 | 2.41 | 2.47 | 2.15 | 1.95 | 2.16 |
| No. reflections | 32287 | 53439 | 46881 | 25245 | 24604 | 24637 | 33342 | 49065 | 32566 |
| $R_{work}/R_{free}$ (%) | 22.6/29.5 | 18.2/22.1 | 23.0/25.7 | 19.1/24.8 | 17.8/23.3 | 20.0/25.7 | 18.1/23.3 | 17.8/21.4 | 17.4/22.7 |
| No. atoms | | | | | | | | | |
| Protein | 10709 | 5245 | 5242 | 5102 | 5237 | 5137 | 5270 | 5254 | 5247 |
| Ligand/ion/water | –/–/– | –/–/714 | –/2/556 | –/1/136 | –/1/176 | –/–/115 | 14/2/357 | 16/2/525 | 34/2/379 |
| B-factors ($Å^2$) | | | | | | | | | |
| Protein | 137.1 | 17.9 | 21.2 | 46.4 | 36.1 | 46.0 | 24.7 | 28.7 | 21.3 |
| Ligand/ion/water | –/–/– | –/–/25.0 | –/32.4/26.3 | –/86.6/40.5 | –/57.0/35.8 | –/–/41.1 | 38.1/25.6/26.8 | 42.0/27.6/34.0 | 35.9/23.7/24.8 |
| RMSD | | | | | | | | | |
| Bond lengths (Å) | 0.013 | 0.008 | 0.008 | 0.008 | 0.008 | 0.009 | 0.007 | 0.008 | 0.008 |
| Bond angles (°) | 1.619 | 1.080 | 1.040 | 1.136 | 1.011 | 1.158 | 0.969 | 0.959 | 1.004 |
| Ramachandran statistics [b] | 96.7/3.1/0.2 | 97.8/1.9/0.3 | 98.3/1.5/0.2 | 97.2/2.6/0.3 | 97.6/2.2/0.2 | 95.8/4.1/0.2 | 97.6/2.2/0.2 | 98.5/1.3/0.2 | 98.4/1.5/0.2 |

[a] Values in parentheses are for highest-resolution shell.

[b] These values are expressed in percentages for “favored, allowed, and disallowed” regions in Ramachandran plots.

**Table S2. Representative viral EN sequences from five human-infecting *Bunyavirales* families and influenza viruses for phylogenetic analysis.**

| Family | Organism | UniProt ID | Full-length L protein (amino acids) | Sequence boundaries of the EN domain |
|---|---|---|---|---|
| *Peribunyaviridae* | Oropouche virus | A0A0D3R3W3 | 2252 | 1-188 |
| *Peribunyaviridae* | Akabane virus | A0A0M5L4E9 | 2251 | 1-188 |
| *Peribunyaviridae* | Bovine Schmallenberg virus (isolate Bovine/BH80/Germany/2011) | H2AM11 | 2254 | 1-188 |
| *Peribunyaviridae* | Bunyamwera virus | P20470 | 2238 | 1-190 |
| *Peribunyaviridae* | Bunyavirus La Crosse | Q38PK9 | 2263 | 1-190 |
| *Peribunyaviridae* | Ngari virus | R4UYG9 | 2238 | 1-190 |
| *Hantaviridae* | Imjin virus | A0A075IK44 | 2149 | 1-190 |
| *Hantaviridae* | Thottapalayam virus | A0A075IM71 | 2150 | 1-190 |
| *Hantaviridae* | Orthohantavirus sangassouense | H6WCR0 | 2151 | 1-190 |
| *Hantaviridae* | Puumala virus (strain Sotkamo/V-2969/81) | P0C760 | 2156 | 1-190 |
| *Hantaviridae* | Hantaan virus (strain 76-118) (Korean hemorrhagic fever virus) | P23456 | 2151 | 1-190 |
| *Hantaviridae* | Seoul virus (strain 80-39) | P27314 | 2151 | 1-190 |
| *Hantaviridae* | Dobrava-Belgrade orthohantavirus (Dobrava virus) | Q806Y6 | 2151 | 1-190 |
| *Hantaviridae* | Sin Nombre orthohantavirus (Sin Nombre virus) | Q89709 | 2153 | 1-190 |
| *Hantaviridae* | Andes orthohantavirus (Andes virus) | Q9E005 | 2153 | 1-190 |
| *Hantaviridae* | Tula orthohantavirus (Tula virus) | Q9YQR5 | 2153 | 1-190 |
| *Phenuiviridae* | Sandfly fever sicilian virus | A0A096ZSN8 | 2090 | 1-223 |
| *Phenuiviridae* | Rift valley fever virus | A2SZS3 | 2092 | 1-224 |
| *Phenuiviridae* | sandfly fever Turkey virus | D9YRN2 | 2090 | 1-223 |
| *Phenuiviridae* | SFTS phlebovirus (isolate SFTSV/Human/China/HB29/2010) | F1BA46 | 2084 | 1-230 |
| *Phenuiviridae* | Sandfly fever Naples virus | I1T351 | 2095 | 1-226 |
| *Phenuiviridae* | Heartland virus | J3TRD1 | 2084 | 1-230 |
| *Phenuiviridae* | Uukuniemi virus (strain S23) | P33453 | 2103 | 1-225 |
| *Phenuiviridae* | Toscana virus | P37800 | 2095 | 1-226 |
| *Arenaviridae* | CAS virus | J7HBG8 | 2046 | 1-179 |
| *Arenaviridae* | Lassa virus (strain Mouse/Sierra Leone/Josiah/1976) | O09705 | 2218 | 1-180 |
| *Arenaviridae* | Lymphocytic choriomeningitis virus (strain Armstrong) | P14240 | 2210 | 1-180 |
| *Arenaviridae* | Tacaribe virus (strain Franze-Fernandez) (TCRV) | P20430 | 2210 | 1-181 |

| | | | | |
|---|---|---|---|---|
| *Arenaviridae* | Junin mammarenavirus | Q0ZEL1 | 2210 | 1-181 |
| *Arenaviridae* | Ippy mammarenavirus (isolate Rat/Central African Republic/Dak An B 188 d/1970) | Q27YE1 | 2208 | 1-180 |
| *Arenaviridae* | Mobala mammarenavirus (isolate Rat/Central African Republic/Acar 3080/1983) | Q27YE5 | 2220 | 1-180 |
| *Arenaviridae* | Machupo virus | Q6IUF8 | 2209 | 1-181 |
| *Nairoviridae* | Xinzhou Spider Virus | A0A0B5KTX6 | 4029 | 440-728 |
| *Nairoviridae* | Issyk-Kul virus | A0A0M4KEL9 | 3992 | 606-893 |
| *Nairoviridae* | Tofla virus | A0A125T1F0 | 3950 | 577-904 |
| *Nairoviridae* | Orthonairovirus bushkeyense | A0A191KWB0 | 3981 | 647-928 |
| *Nairoviridae* | Tamdy virus | A0A5P9K616 | 3980 | 592-909 |
| *Nairoviridae* | Kasokero virus | A0A7S6HEL7 | 3972 | 611-904 |
| *Nairoviridae* | Yezo virus | A0A9E9C2C9 | 3938 | 600-878 |
| *Nairoviridae* | Hazara virus | A6XA53 | 3923 | 577-879 |
| *Nairoviridae* | Nairobi sheep disease virus | D0PRM7 | 3991 | 580-943 |
| *Nairoviridae* | Erve virus | J3RTH4 | 3863 | 548-830 |
| *Nairoviridae* | Dugbe virus (isolate ArD44313) | Q66431 | 4036 | 584-989 |
| *Nairoviridae* | Crimean-Congo hemorrhagic fever virus (strain Nigeria/IbAr10200/1970) | Q6TQR6 | 3945 | 585-898 |
| *Orthomyxoviridae* | Influenza A virus (strain A/Korea/426/1968 H2N2) | P13170 | 716 | 1-210 |
| *Orthomyxoviridae* | Influenza A virus (strain A/Wilson-Smith/1933 H1N1) | P15659 | 716 | 1-210 |
| *Orthomyxoviridae* | Influenza A virus (strain A/Hong Kong/1/1968 H3N2) | Q91MA9 | 716 | 1-210 |
| *Orthomyxoviridae* | Influenza B virus (B/Victoria/02/1987) | A4D5P7 | 726 | 1-207 |
| *Orthomyxoviridae* | Influenza B virus (strain B/Yamagata/16/1988) | A4D5Q8 | 726 | 1-207 |
| *Orthomyxoviridae* | Influenza C virus (strain C/JJ/1950) | P13878 | 709 | 1-190 |
| *Orthomyxoviridae* | Influenza C virus (strain C/Johannesburg/1/1966) | Q9IMP5 | 709 | 1-190 |

### Table S3. The EN sequences of *Nairoviridae* family for evolutionary conservation analysis.

| Genus | Organism | UniProt ID | Full-length L protein (amino acids) | Sequence boundaries of the EN domain |
|---|---|---|---|---|
| *Orthonairovirus* | Nairobi sheep disease virus | A0A0A7H7L5 | 3991 | 579-943 |
| *Orthonairovirus* | Wenzhou Tick Virus | A0A0B5KKH6 | 3967 | 593-900 |
| *Orthonairovirus* | Huangpi Tick Virus 1 | A0A0B5KTT9 | 3914 | 596-877 |
| *Orthonairovirus* | Leopards Hill virus | A0A0B6VH63 | 3967 | 611-899 |
| *Orthonairovirus* | Issyk-Kul virus | A0A0M4KEL9 | 3992 | 605-893 |
| *Orthonairovirus* | Keterah virus | A0A0M4KPD1 | 3992 | 605-893 |
| *Orthonairovirus* | Orthonairovirus thiaforaense | A0A0M4KY89 | 3861 | 551-833 |
| *Orthonairovirus* | Yogue virus | A0A0M4L956 | 3965 | 609-900 |
| *Orthonairovirus* | Gossas virus | A0A0M5KTS8 | 3991 | 606-890 |
| *Orthonairovirus* | Tofla virus | A0A0U5BRX5 | 3511 | 576-904 |
| *Orthonairovirus* | Taggert virus | A0A142J8F6 | 3906 | 589-869 |
| *Orthonairovirus* | Abu Hammad virus | A0A191KW45 | 3955 | 595-878 |
| *Orthonairovirus* | Abu Mina virus | A0A191KW46 | 3974 | 597-880 |
| *Orthonairovirus* | Bandia virus | A0A191KW77 | 3967 | 605-886 |
| *Orthonairovirus* | Clo Mor virus | A0A191KW84 | 3888 | 586-866 |
| *Orthonairovirus* | Great Saltee virus | A0A191KW99 | 3968 | 628-910 |
| *Orthonairovirus* | Punta Salinas virus | A0A191KWB2 | 3986 | 653-935 |
| *Orthonairovirus* | Sapphire II virus | A0A191KWC5 | 3956 | 590-873 |
| *Orthonairovirus* | Soldado virus | A0A191KWD1 | 3957 | 624-906 |
| *Orthonairovirus* | Tunis virus | A0A191KWD6 | 3955 | 595-878 |
| *Orthonairovirus* | Zirqa virus | A0A191KWE7 | 3986 | 642-924 |
| *Orthonairovirus* | Orthonairovirus chimense | A0A1S5NQC4 | 4010 | 612-895 |
| *Orthonairovirus* | Geran virus | A0A1S5NQY1 | 3969 | 605-884 |
| *Orthonairovirus* | Orthonairovirus artashatense | A0A1S5NU79 | 3951 | 580-861 |
| *Orthonairovirus* | Burana virus | A0A1S5NVP2 | 3972 | 593-903 |
| *Orthonairovirus* | Vinegar Hill virus | A0A2H4X1S4 | 3948 | 593-876 |
| *Orthonairovirus* | Pacific coast tick nairovirus | A0A2R2WU07 | 4061 | 592-986 |
| *Orthonairovirus* | Estero Real virus | A0A346JIY0 | 3925 | 594-874 |
| *Orthonairovirus* | Tacheng Tick Virus 1 | A0A5C0C9B6 | 3962 | 587-905 |

| | | | | |
|---|---|---|---|---|
| *Orthonairovirus* | Tamdy virus | A0A5P9K616 | 3980 | 591-909 |
| *Orthonairovirus* | Meram virus | A0A7G7Y1S5 | 4007 | 576-961 |
| *Orthonairovirus* | Camel Crimean-Congo hemorrhagic fever orthonairovirus | A0A7S6HGL1 | 3945 | 584-898 |
| *Orthonairovirus* | Kasokero virus | A0A7S6KQ42 | 3972 | 610-904 |
| *Orthonairovirus* | Songling virus | A0A7U3T2V7 | 3951 | 592-888 |
| *Orthonairovirus* | Avalon virus | A0A859D691 | 3903 | 587-867 |
| *Orthonairovirus* | Sulina virus | A0A893CDI9 | 3940 | 599-878 |
| *Orthonairovirus* | Wufeng Crocidura attenuata orthonairovirus 1 | A0A8T9KN41 | 3919 | 579-861 |
| *Orthonairovirus* | Yezo virus | A0A9E9C2C9 | 3938 | 599-878 |
| *Orthonairovirus* | Cencurut virus | A0A9Y1HTN0 | 3887 | 564-847 |
| *Orthonairovirus* | Meihua Mountain virus | A0AAE9HT59 | 3923 | 576-878 |
| *Orthonairovirus* | Pangolin orthonairovirus | A0AAE9RUZ8 | 3970 | 593-900 |
| *Orthonairovirus* | Hazara virus | A6XA53 | 3923 | 576-879 |
| *Orthonairovirus* | Kupe virus | B8PWH5 | 4050 | 583-1003 |
| *Orthonairovirus* | Erve virus | J3RTH4 | 3863 | 547-830 |
| *Orthonairovirus* | Dugbe virus (isolate ArD44313) (DUGV) | Q66431 | 4036 | 583-989 |
| *Orthonairovirus* | Crimean-Congo hemorrhagic fever virus (strain Nigeria/IbAr10200/1970) (CCHFV) | Q6TQR6 | 3945 | 584-898 |
| *Sabavirus* | South Bay virus | A0A076E6R2 | 4536 | 1081-1397 |
| *Striwavirus* | Sanxia Water Strider Virus 1 | A0A0B5KF82 | 3934 | 345-640 |
| *Shaspivirus* | Shayang Spider Virus 1 | A0A0B5KRX1 | 4403 | 592-976 |
| *Xinspivirus* | Xinzhou Spider Virus | A0A0B5KTX6 | 4029 | 439-728 |
| *Norwavirus* | Grotenhout virus | A0A1V0EFI9 | 4812 | 1349-1665 |
| *Ocetevirus* | red goblin roach virus 1 | A0A7D7JJ32 | 4015 | 347-632 |
